# Proteomic Insights into Strong and Weak Biofilm Formation in *Acinetobacter baumannii* for Potential Therapeutic Targets

**DOI:** 10.1101/2025.10.02.680171

**Authors:** Umarani Brahma, Akash Suresh, Aafreen Kamila, Usha S, Arun Kumar S.V, Harshala Baddi, Siva Singothu, Paresh Sharma, Vasundhra Bhandari

## Abstract

*Acinetobacter baumannii* is notable for its biofilm-forming abilities, which aid in its tolerance to antibiotics, adding to antimicrobial resistance. The clinical isolates present varied biofilm-forming capacity; hence, understanding the molecular determinants that result in strong biofilm development is crucial for drug target identification. This study is the first of its kind to compare proteome profiling of strong and weak biofilm-forming *A. baumannii* clinical isolates. Comparative proteomic profiling revealed 42 differentially regulated proteins. It was observed that in strong biofilm forming isolate NlpA, uL16, DNA gyrase B, acetyl-CoA carboxylase, and purl etc. were upregulated highlighting a dynamic reprogramming of cellular functions that promotes biofilm formation, stress adaptation, and immune evasion. In contrast, EF-Tu, ribosome hibernation factors, and T6SS components were downregulated, suggesting a lack of biosynthesis and stress adaptability. These findings suggest a metabolic downshift and a possible energy conservation mechanism under conditions less favorable for strong biofilm development. Additionally, several uncharacterized proteins were identified, highlighting potential novel factors in biofilm regulation and virulence that warrant further investigation. The proteomics data correlated with qPCR findings, providing support for the unknown regulators of biofilm formation that were identified in this study. Key proteins such as *nlpA*, 6,7-dimethyl-8-ribityllumazine synthase and DNA gyrase B emerged as potential therapeutic targets.

Graphical Abstract

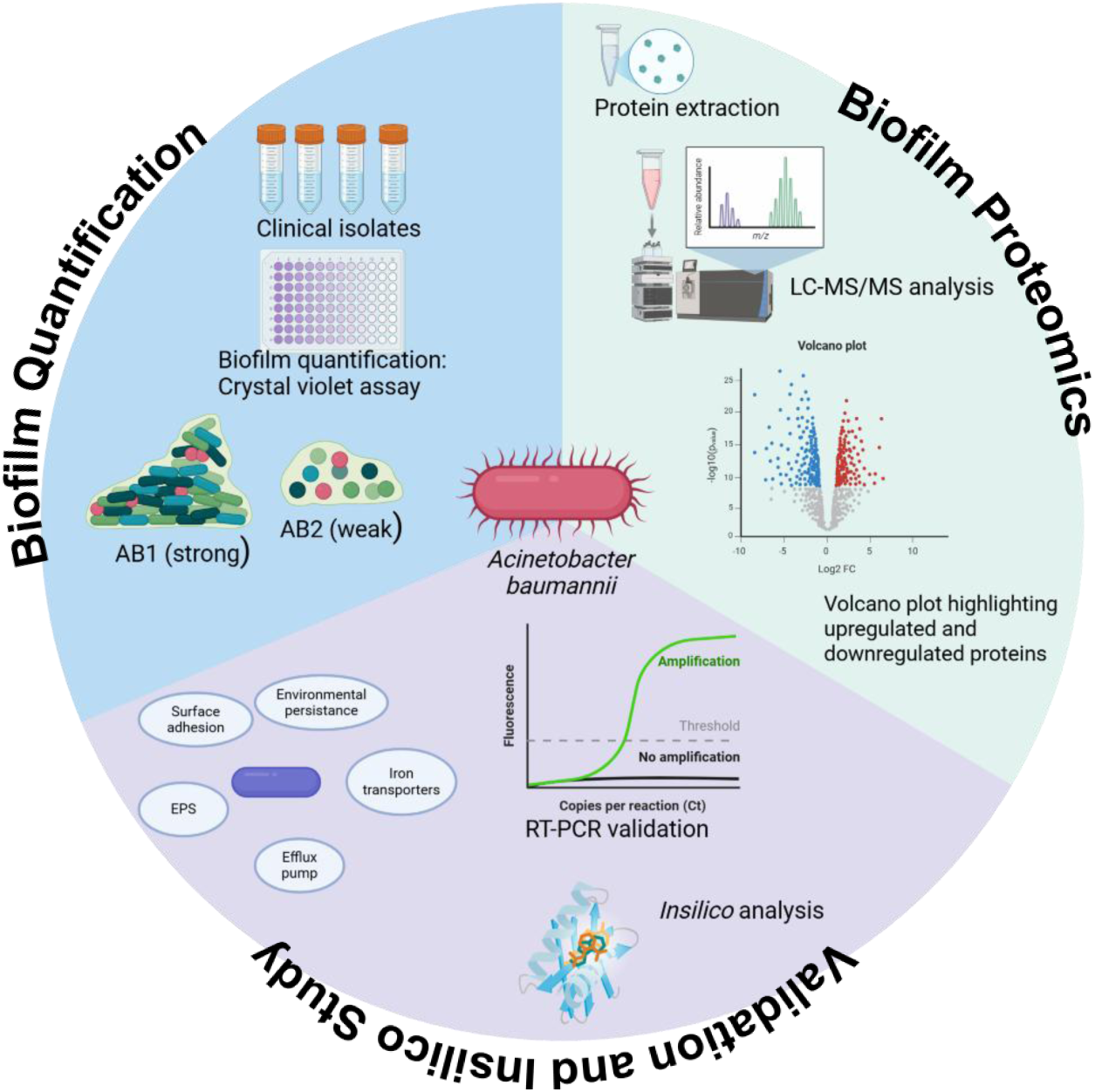

## 1. Introduction

Antimicrobial resistance (AMR) is an ascending global health crisis, and *Acinetobacter baumannii* stands out as one of the most daunting threats due to its exceptional ability to resist antibiotics and persist in hospital environments ^1^. This Gram-negative opportunistic pathogen is a notable member of the “ESKAPE” group of bacteria, known for evading the effects of antimicrobial agents ^2^. *A. baumannii* poses a significant therapeutic challenge, despite the development of new antibiotics and adjuvants ^3^. The World Health Organization (WHO) and the Centers for Disease Control and Prevention (CDC) have classified it as a critical priority pathogen, underscoring the urgent need for novel therapeutic strategies ^1^. Also reported by India’s AMR surveillance network, a staggering 80% of *A. baumannii* isolates exhibited resistance to imipenem, bringing attention to a critical threat in the fight against drug-resistant infections ^4^. Due to its dynamic genome, the isolates are different from one another and make it hard to define them as a single species ^1^.

Another feather is its robust capacity to form biofilms, a structured microbial community encased in a self-produced extracellular matrix ^5^. Their biofilm-forming ability confers them remarkable survival advantages, which include resistance to desiccation, disinfectants, oxidative stress, and host immune responses. Biofilm formation on medical devices such as catheters, ventilators, and prosthetics facilitates persistent infections and complicates treatment, often leading to chronic and recurrent infections ^6^. Within biofilms, bacteria exhibit up to 1,000-fold increased resistance to antibiotics compared to their planktonic counterparts, rendering conventional therapies largely ineffective ^7^.

The biofilm lifestyle of *A. baumannii* is intricately linked to its pathogenicity ^8^. It contributes to the persistence of infections such as ventilator-associated pneumonia, catheter-associated urinary tract infections, and bloodstream infections, particularly in intensive care settings ^9^. These infections are associated with prolonged hospital stays, increased healthcare costs, and elevated mortality rates ^10^. The bacterium’s ability to survive on abiotic surfaces and evade host defenses through biofilm-mediated dormancy and immune modulation makes it a formidable nosocomial pathogen ^11^. Biofilm formation in *A. baumannii* is regulated by a complex network of genetic and environmental factors, including the csuA/BABCDE operon, quorum-sensing systems, and surface-associated proteins such as BfmS, chaperone-usher pili, outer membrane protein A and several others. These elements orchestrate adhesion, maturation, and maintenance of the biofilm architecture ^5, 12^. Clinical isolates often exhibit enhanced biofilm-forming capabilities compared to environmental strains, further complicating infection control and treatment efforts ^13^.

Despite the critical role of biofilms in *A. baumannii* pathogenesis, most drug discovery efforts have historically focused on its planktonic phenotype ^14^. Recent advances in proteomics have opened new avenues for understanding the molecular underpinnings of biofilm formation ^15^. This study employs a label-free global proteomic approach to compare the proteomes of strong and weak biofilm-forming multidrug-resistant (MDR) clinical isolates of *A. baumannii*. By elucidating differential protein expression profiles, this work aims to uncover biofilm-associated regulators and pathways, offering insights into potential therapeutic targets to combat biofilm-mediated infections.

## 2. Experimental Section

### 2.1 Sample collection and Molecular characterization of *A. baumannii* clinical isolates

The current study used 25 clinical isolates obtained from patients with blood-borne, respiratory tract, and skin infections reporting to ESIC Hospital, Hyderabad, India. The cultures were maintained in TSB (Tryptone Soy Broth, Himedia, India) medium at 37 °C for 16 to 20 hours. Species confirmation was carried out through universal 16S rRNA gene sequencing. Genomic DNA was isolated using a Wizard genomic kit (Promega, Madison, WI, USA) with slight modifications. In brief, 3 mL of the culture was centrifuged for 10 minutes at 4000× g to form a pellet. The pellet was washed twice with 1x PBS and was thereafter resuspended in 500 µL of 50 mM Tris-EDTA containing 100 µl of 20mg/mL lysozyme for two hours at 37 °C before processing as per the manufacturer’s guidelines. The DNA was diluted in sterile deionized water, and its purity was assessed using a Nanodrop (Thermo Scientific, Waltham, MA, USA). The 16S rRNA gene was amplified as previously reported ^16^ and then sequenced using Sanger sequencing at Ira Biotech, Hyderabad, India. A quality control strain, ATCC 19606, was used in studies of biofilm and antimicrobial susceptibility, in accordance with the Clinical and Laboratory Standards Institute (CLSI) guidelines ^17^.

### 2.2 Antimicrobial susceptibility studies

Antimicrobial susceptibility testing was performed using both disk diffusion and microbroth dilution methods in *A. baumannii* clinical isolates (n=25), and ATCC 19606 was used as a control strain as per CLSI guidelines. The antibiotics tested for susceptibility included carbapenems (imipenem, meropenem), fluoroquinolones (ciprofloxacin), tetracyclines (minocycline, doxycycline), cephalosporins (ceftazidime, ceftriaxone), lipopeptides (colistin), and the β-lactam combination drug piperacillin-tazobactam (Himedia, India). Microbroth dilution experiment with resazurin dye (Sigma, Bangalore, India) as an indicator was used to evaluate the susceptibility against imipenem, colistin, ciprofloxacin, minocycline, and ceftazidime ^18^. Disk diffusion assay was performed against ceftriaxone (30 µg), doxycycline (30 µg), and piperacillin-tazobactam (2 µg). Isolates were categorized according to resistance profiles as extensively drug-resistant (XDR; susceptible to one or two classes), Pan-drug resistant ^19^.

### 2.3 Biofilm formation

The biofilm formation was determined using crystal violet (CV) (Sigma, Bangalore, India) assay with slight modifications to the standard protocol ^20^. In brief, 200 µL of an overnight culture grown in TSB + 2% glucose (1:200 dilution) was added into each well of a 96-well microtiter plate and incubated at 37°C for 24 h under static conditions. Subsequent to incubation, wells were washed thrice with 1× PBS and fixed with 200 µL of methanol for 15 minutes. The plates were air-dried for 20 minutes, followed by the addition of 100 µL of 0.2% CV solution and incubated at room temperature for 15 minutes. Excess dye was removed by washing with distilled water, and the adhered stain was solubilized using 100 µL of 33% glacial acetic acid. The biomass of the biofilm was assessed by measuring absorbance at 590 nm using a multimode plate reader (Cytation 5). The biofilm-forming capacity of isolates was categorized according to optical density (OD) values as follows: non-adherent (ODs ≤ ODc), weak (ODc < ODs ≤ 2×ODc), moderate (2×ODc < ODs ≤ 4×ODc), and strong (ODs > 4×ODc), where ODc denotes the mean OD of the negative control and ODs signifies the OD of the examined strains ^14^.

### 2.4 Protein extraction from *A. baumannii* biofilms

The biofilms were grown as described previously ^14, 21^. Briefly, overnight cultures of the clinical isolates were grown in TSB with 2% glucose at 37°C overnight, and 10^7^ cells/ml were then seeded into 6-well flat-bottomed polystyrene microtiter plates and incubated at 37 °C for 48 h under static conditions to facilitate biofilm formation. The supernatant was removed, and the plates were washed twice with 1x PBS to remove any unadhered cells. Biofilm formation for each of the two isolates was performed in thrice in triplicate. Biofilm cells from each culture were scraped, pooled, and processed separately, followed by centrifugation at 4000 xg for 10 minutes at 4°C and then washed twice with 1x PBS before protein extraction. The resultant pellet was then resuspended in 1x TE buffer, and 20 mg/mL lysozyme was added and incubated for 1 h at 37°C before adding RIPA lysis buffer (HiMedia, India) for isolating protein. The cells were mixed thoroughly, followed by sonication on ice for 5 min at a 10 sec ON/OFF pulse. The homogenized cells were centrifuged at 16,000 g for 10 min at 4 °C to obtain the protein in the supernatant. Protein estimation was carried out using the BCA assay (Pierce - Thermo-Scientific).

### 2.5 Sample preparation for quantitative label-free proteomics

100µg of protein was processed for LC-MS/MS analysis using the in-solution trypsin digestion method. To the protein 1% sodium deoxycholate was added, followed by reduction with 20 mM DTT at 57°C for 1 h. Next, alkylation was carried out with 200 mM iodoacetamide (IAA) for 1 h at room temperature in the dark. The proteins were then digested with trypsin-protease (Thermo-scientific) (1:100 wt/wt) overnight at 37°C. The digestion reaction was stopped by adding 0.1% formic acid. The digested peptides were then purified using C_18_ spin columns. The purified peptides were concentrated using vacuum evaporator and finally resuspended in 0.1% trifluoroacetic acid ^22^.

### 2.6 Peptide mixtures were subjected to mass spectrometric analysis

The peptides were analyzed with a Q Exactive HF-Orbitrap mass spectrometer (Thermo Fisher Scientific) coupled with an Ultimate 3000 RSLCnano LC system (Thermo Fisher Scientific). The peptides were injected into a reverse-phase C18 column (PepMap RSLC C18, 2 μm, 100 Å, 75 μm by 50 cm; Thermo Fisher Scientific) and separated by a gradient flow of solvent B (0.1% formic acid in 80/20 acetonitrile/water) from 5% to 90% for 60 min. A parent ion scan was performed with a scan range of 375 to 1,600 *m/z* with a resolution of 60,000. The top 25 intense peaks were fragmented by higher energy collision-induced dissociation (HCD) fragmentation in MS/MS with a resolution of 15,000 ^22^.

### 2.7 Data Processing

The experimental design consisted of two biological groups: AB1 (strong biofilm former) and AB2 (weak biofilm former), each comprising two biological replicates. Label-Free Quantitative (LFQ) proteomic analysis was performed using Perseus-MaxQuant software. Initially, RAW files were obtained from a Q Exactive HF-Orbitrap mass spectrometer and analyzed using MaxQuant (version 2.7.3). *A. baumannii* protein sequences were obtained from UniProt (version 2025_01). Within MaxQuant, “matching between runs” and “LFQ” were chosen with the default value settings. Downstream analysis was carried out in the Perseus (version 2.1.4). The ‘proteinGroups.txt’ output file obtained from MaxQuant was imported into the Perseus suite, and the pertinent columns were chosen for both strong and weak biofilm-forming strains and subsequently log transformed. Quantitative profiles underwent a filtration process to address missing values, with independent filtering applied to each strain, ensuring that only proteins quantified in both replicates were retained. Imputation of missing values was conducted (width 0.3, down shift 1.8) before the integration of the tables and the execution of the multi-volcano analysis. The s0 and FDR parameters for the multi-volcano analysis were selected based on visual inspection for the strong biofilm strain (higher confidence, s0 = 1, FDR = 0.01%) and the weak biofilm strain (lower confidence, s0 = 1, FDR = 0.2%), to minimize the number of significantly depleted proteins across all experiments ^23^.

### 2.8 Experimental Design and Statistical Rationale

The protein expression profiles of strains AB1 and AB2 served as the foundation for identifying the patterns of up-regulation and down-regulation. Values that were missing were filled in using the normal distribution method. Abundance values underwent log₂ normalization, subsequently followed by Z-score standardization. The Z-score values underwent additional processing through Student’s t-test to determine the significance of the proteins. Significance was calculated using Benjamini-Hochberg, with a 0.05 FDR established as the significance cutoff. The correlation plot, utilizing Pearson’s correlation coefficient, was employed to visualize the relationship between each sample. Principal component analysis (PCA) was conducted to assess variations among biological replicates and within biological replicates. Only the proteins that exhibited up-regulation or down-regulation with both AB1 and AB2 were selected for subsequent gene ontology and pathway enrichment analysis. The proteins exhibiting analogous expression patterns, in conjunction with cellular components and biological functions, were depicted through a heat map visualization. Four biological replicates were utilized for the real-time PCR analysis. The results’ significance was examined through the analysis of variance. Differences were deemed significant when p-values fell below 0.05 ^21^ ^14^.

### 2.9 Gene Ontology Analysis

The differentially expressed proteins have been studied through the STRING database (version 12.0), (https://string-db.org) to anticipate the functional interactions between proteins. The analysis was performed using a confidence score threshold of > 0.9, focusing exclusively on experimentally validated and database-derived interactions. The interaction networks generated were visualized and subjected to further analysis to pinpoint essential hubs and enriched pathways.

### 2.10 RNA Extraction and Real-time PCR

RNA was extracted from 48 h-old *A. baumannii* biofilms using the Macherey Nagel RNA isolation kit (NucleoSpin® RNA, 740955.10) following the manufacturer’s instructions. Subsequently, cDNA was synthesized from approximately 2.5 µg of RNA using the Clontech cDNA synthesis kit (PrimeScript™ 1st strand cDNA Synthesis Kit, 6110B). The 2^−ΔΔCₜ^ method was used to calculate the relative gene expression using SYBR green dye and was carried out on the QuantStudio 7 Pro real-time detection equipment (Applied Biosystems). Table 1 presents the list of primers utilized in this study.

**Table 1:**
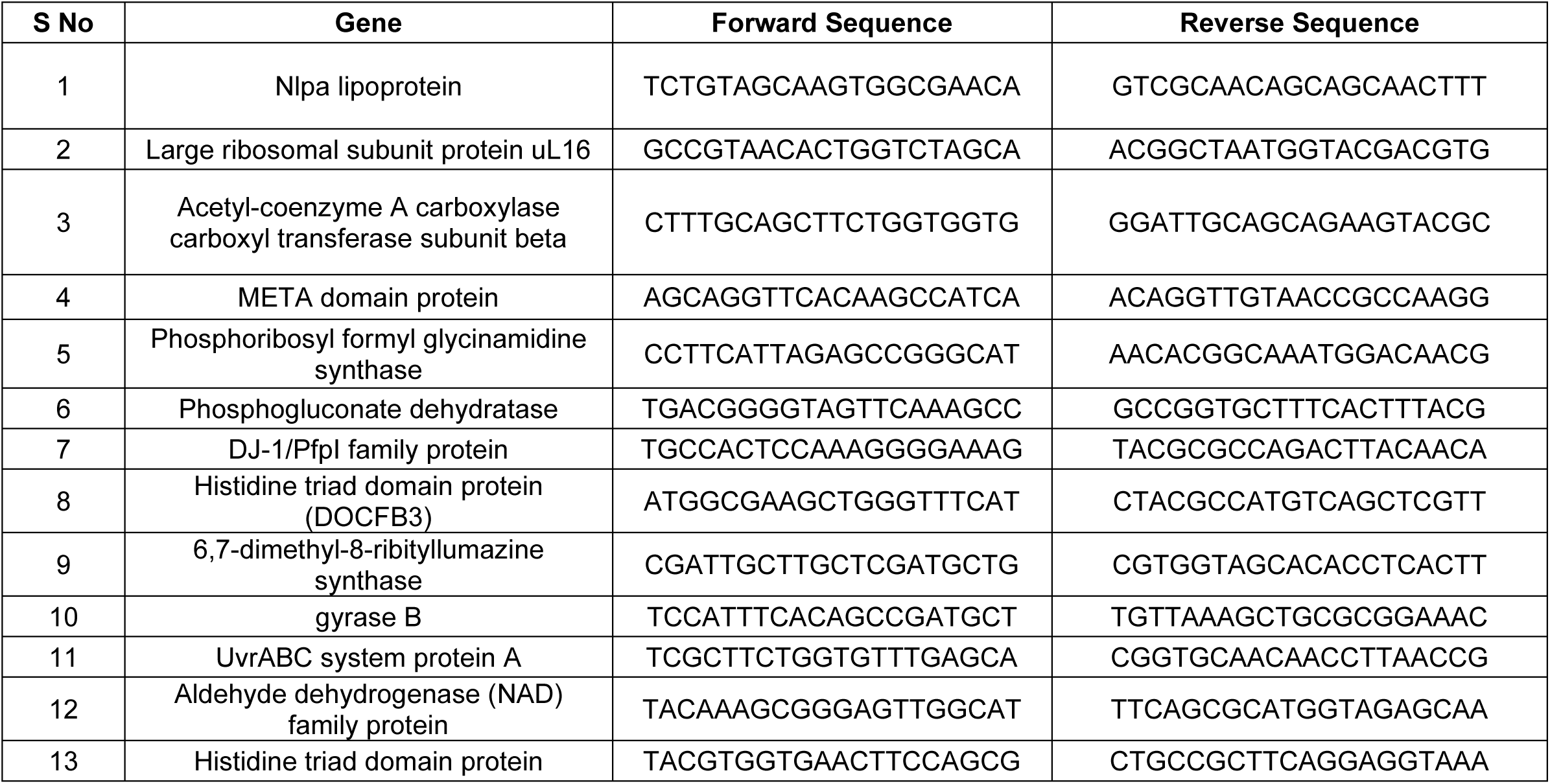
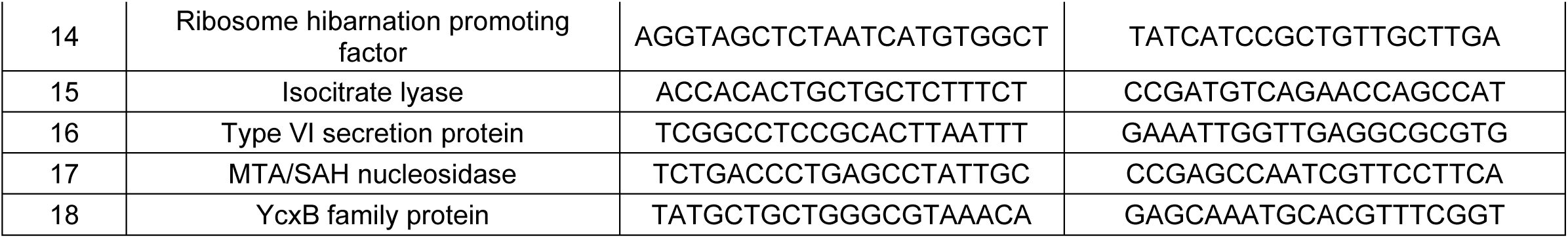
List of primers used in the study.

## 3. Results and Discussion

### 3.1 Antimicrobial and biofilm profiling of *A.baumannii* clinical isolates

In the study, 25 confirmed 16S rRNA gene-sequenced *A. baumannii* clinical isolates were included. These isolates were subjected to antimicrobial susceptibility testing against the commonly used antibiotics using microbroth dilution and disk diffusion assay. The majority of isolates (92%, n = 23), exhibited resistance to carbapenems, cephalosporins, fluoroquinolones, tetracyclines, and β-lactam/β-lactamase inhibitor combination, and were classified as extensively drug-resistant (XDR). Only 8% (n = 2) of isolates were found to be resistant to the lipopeptide (colistin) class of antibiotics. None of the clinical isolates was found to be pan-sensitive, and one clinical isolate was found to be resistant to all the classes of antibiotics categorized as pan-resistant. The biofilm-forming capacity was assessed in all the isolates using the CV assay, and they were characterized as strong (n = 13; 52%), moderate (n = 8; 25%) and weak (n = 3; 12%) biofilm forming isolates (Figure 1a and 1b).

**Figure 1A:** Antimicrobial resistance profile of *A.baumannii* clinical isolates. Antimicrobial resistance profile of *A.baumannii* clinical isolates. The susceptibility profile of all clinical isolates was determined using microbroth dilution and disc diffusion methods.

**Figure 1B:**
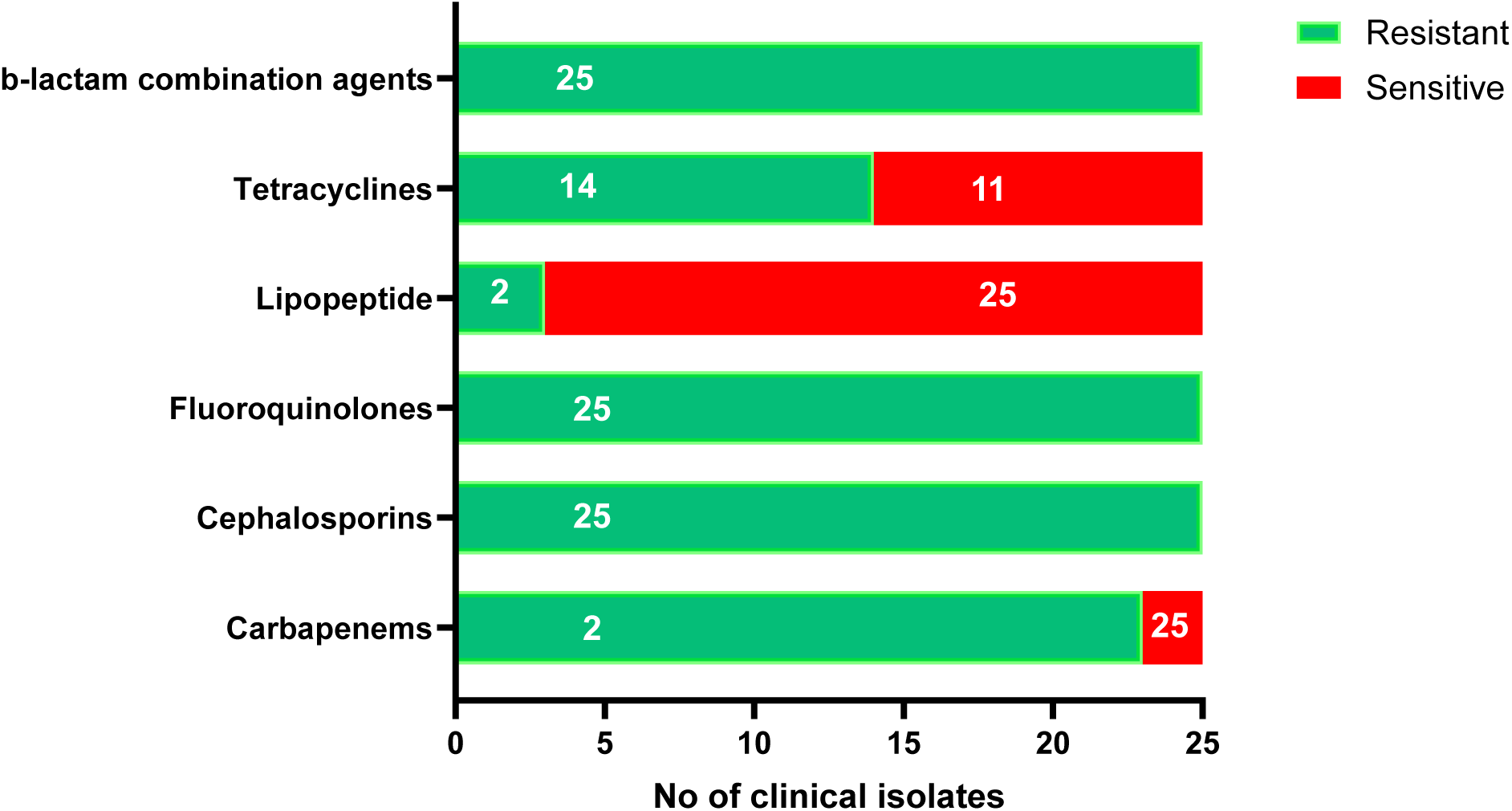
Distribution of biofilm capacity among clinical isolates of *A.baumannii.* Distribution of biofilm capacity among clinical isolates of A.baumannii. Biofilm biomass was quantified using the crystal violet dye by measuring the absorbance at 570 nm.

The capacity of *A. baumannii* to form biofilms is an important virulence characteristic that contributes to the pathogenicity of the bacteria and is thought to play a significant role in its ability to survive in adverse habitats ^24^. The biofilms enhance the virulence of *A. baumannii* by promoting bacterial survival under adverse environmental conditions, including exposure to disinfectants, antibiotics, and host immune defenses ^25^. Importantly, antibiotic resistance phenotypes have a strong influence on biofilm-forming capacity. Previous studies have demonstrated that MDR strains exhibit greater biofilm thickness and higher expression of biofilm-associated virulence genes compared to drug-sensitive isolates. A significant increase in biofilm-forming, drug-resistant *A. baumannii* has been documented in intensive care units (ICUs), and numerous studies have indicated a strong relationship between antimicrobial resistance and biofilm-forming ability. A study reported that 97.1% of the clinical isolates could form biofilms, in which 4.3% possessed weak biofilm formation ability, while 41.4% and 51.4% were moderate and strong biofilm producers ^26^. This observation aligns with our findings, whereby most isolates exhibited moderate to high biofilm-forming capability, hence reinforcing the established association between drug resistance and increased biofilm development. Another study reported that 58% of the isolates showed strong ability to form biofilm, whereas 42% isolates showed moderate biofilm formation ^27^. There is also a report on strong biofilm formation capacity in *A. baumannii* isolated from burn units. These isolates exhibited significant resistance to antibiotics, including carbapenems, and demonstrated co-production of AmpC and ESBLs ^28^. In contrast, a study reported that the sensitive isolates exhibited strong biofilm formation in the initial phase (< 24 h) relative to the resistant isolates, whereas their biofilm-forming capacity dropped in subsequent phases ^26^.

We found two isolates to be colistin resistant, and both displayed strong biofilm formation. This is in-line with a study where colistin-resistant *A. baumannii* clinical isolates exhibited an enhanced biofilm-forming capacity, suggesting that biofilm production may represent an adaptive response associated with the development of colistin resistance ^25^. Whereas the previous study suggests that the correlation between colistin resistance and biofilm formation is dependent upon the specific resistance mechanism involved. Mutations and modifications to LPS maintain biofilm development, therefore enhancing persistence and transmission in healthcare environments ^29^. These results align with our observation that colistin-resistant *A. baumannii* clinical isolates are capable of maintaining strong biofilm-forming ability. This confirms our results that colistin-resistant *A. baumannii* may maintain significant biofilm-forming capability. Whereas in contrast, one study revealed that the acquisition of colistin resistance in two clinical *A. baumannii* isolates was accompanied by a marked reduction in biofilm-forming ability. The limited available information comes from another study that revealed laboratory-evolved Colᴿ mutants, which also demonstrated less biofilm production relative to their susceptible counterparts ^30^.

### 3.2 Identification of differentially regulated proteins in the Strong and Weak Biofilm-forming *A. baumannii* isolates

In taking note of the observed variability in biofilm formation and the distinct phenotypes linked to XDR and colistin resistance associated with the clinical strains, relevant isolates were carefully selected for the proteomics study. Isolates showing strong and weak formation of biofilm, in addition to colistin-resistant and colistin-susceptible, as well as multidrug-resistant phenotypes, were included. This methodology is meant to elucidate the proteome variations that contribute to biofilm heterogeneity and resistance mechanisms, therefore offering insights into the molecular factors that influence virulence and persistence in *A. baumannii*.

A label-free quantitative mass spectrometric analysis was carried out to identify the regulators linked to the persistence of infection related to the biofilm-forming capability of *A. baumannii*. A total of 1066 proteins were identified in both the strong and weak isolates from the proteomic analysis. An analysis of the fold changes between AB1 and AB2 was conducted to get the differential expression profiles. The student’s t-test was then used on all 712 differentially expressed proteins for identifying specific protein markers linked with biofilm characteristics. We observed protein enrichment in pathways associated with energy metabolism, lipid biosynthesis, oxidative stress adaptation, and transport functions critical for biofilm matrix synthesis and durability in the AB1 isolate. Conversely, in strains exhibiting weak biofilm formation, proteins related to ribosome maintenance, RNA processing, and translation initiation were prevalent, indicating a diminished anabolic profile.

Differential expression thresholds were set at log₂ fold change > 2 and p < 0.05. A total of 41 proteins met the significance criteria, comprising 19 upregulated and 22 downregulated proteins in strong biofilm conditions compared to weak (Figure 2). The obtained fold changes were then filtered using population statistics to get the significant threshold cutoff of up-regulated and down-regulated proteins for each of the comparisons performed in the study. To visualize differential expression trends, a heatmap of the 41 significantly altered proteins was generated using hierarchical clustering based on Euclidean distance and average linkage (Figure 3).

**Figure 2:**
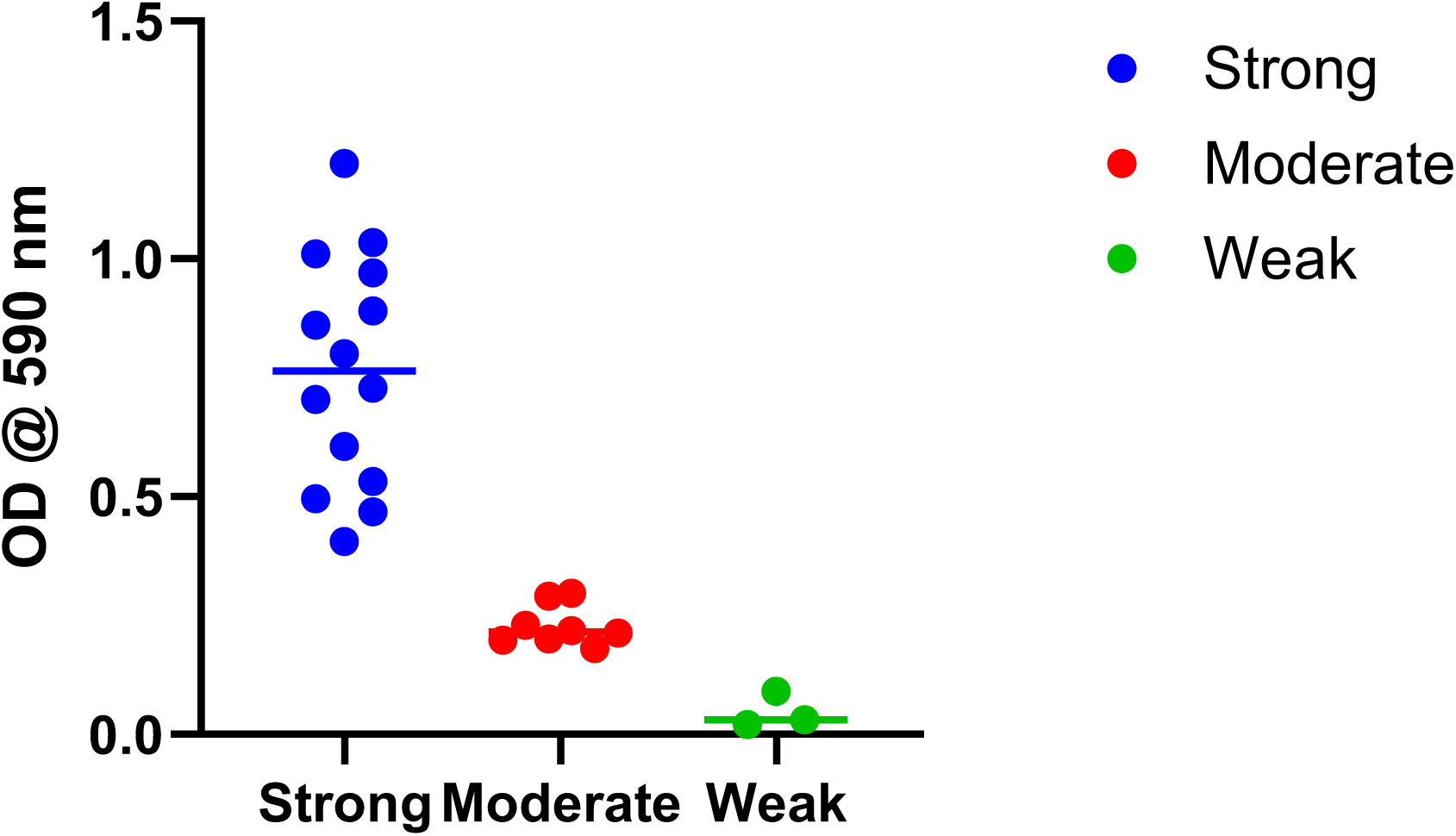
Comparative proteomic heatmap showing upregulated and downregulated proteins in biofilm-forming *A. baumannii*. Protein intensities were scaled and color-coded: red for high expression, green/blue for low expression, and black for intermediate levels. Distinct clustering patterns were observed, separating strong biofilm samples (AB1) from weak biofilm samples (AB2), and grouping proteins based on shared expression dynamics.

**Figure 3:**
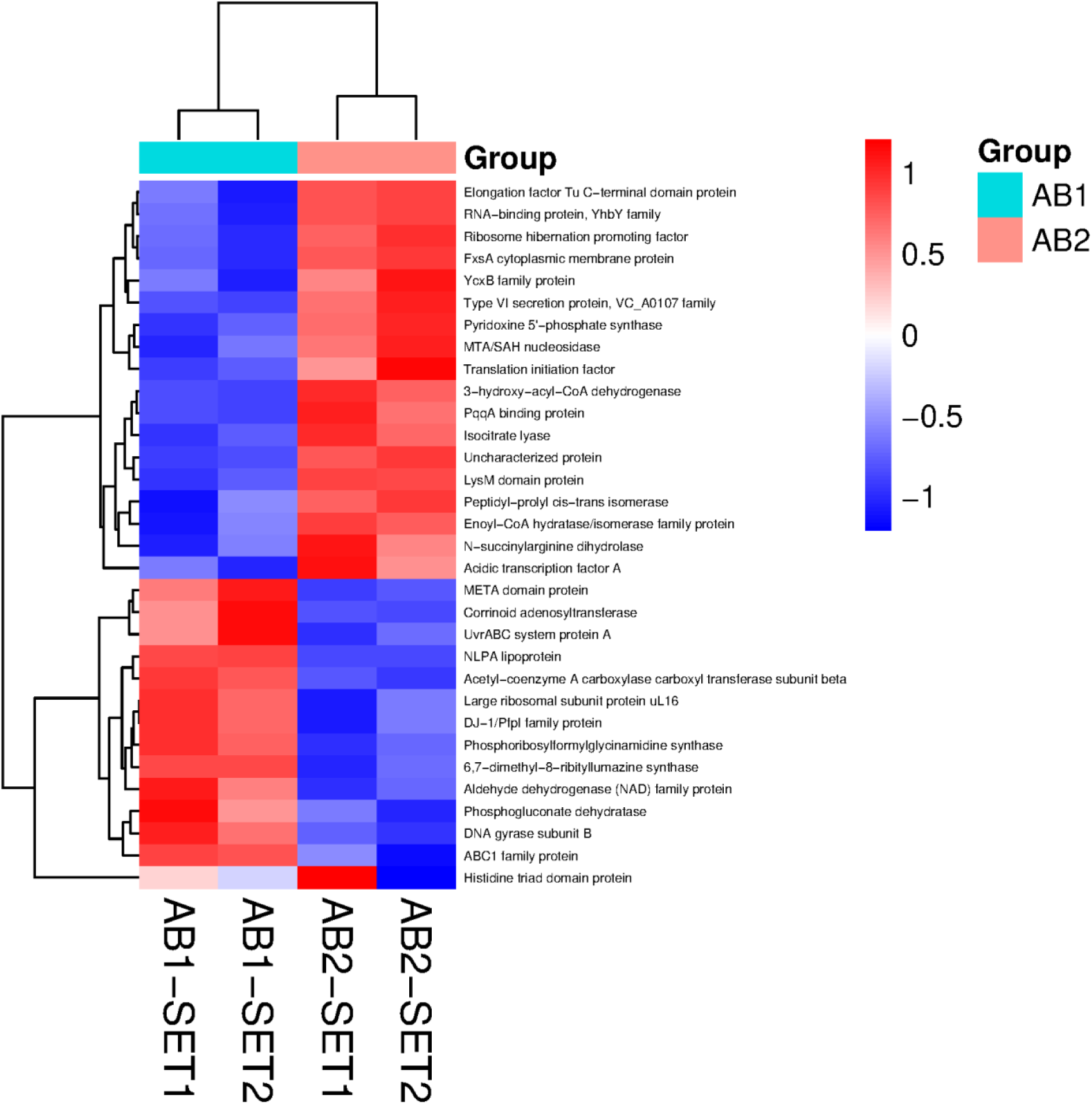
A volcano plot showing significantly differentially expressed proteins between strong and weak biofilm conditions. The x-axis represents fold change (log₂), and the y-axis represents statistical significance (−log₁₀ p value). Proteins significantly upregulated in strong biofilm formers are shown in red, while proteins significantly upregulated in weak biofilm formers are shown in blue. Gray points represent non-significant proteins. Labelled points indicate proteins with the most significant differential expression.

### 3.3 Proteomic remodelling in strong biofilm-forming *A. baumannii*

#### A. Upregulated proteins

The bioinformatic analysis revealed significantly upregulated proteins that could play a vital role in the biofilm formation and antibiotic resistance of *A. baumannii* (Table 2). These proteins include several functional categories, including adhesion, stress adaptation, DNA replication, metabolic reprogramming, and a few uncharacterized proteins.

**Table 2:**
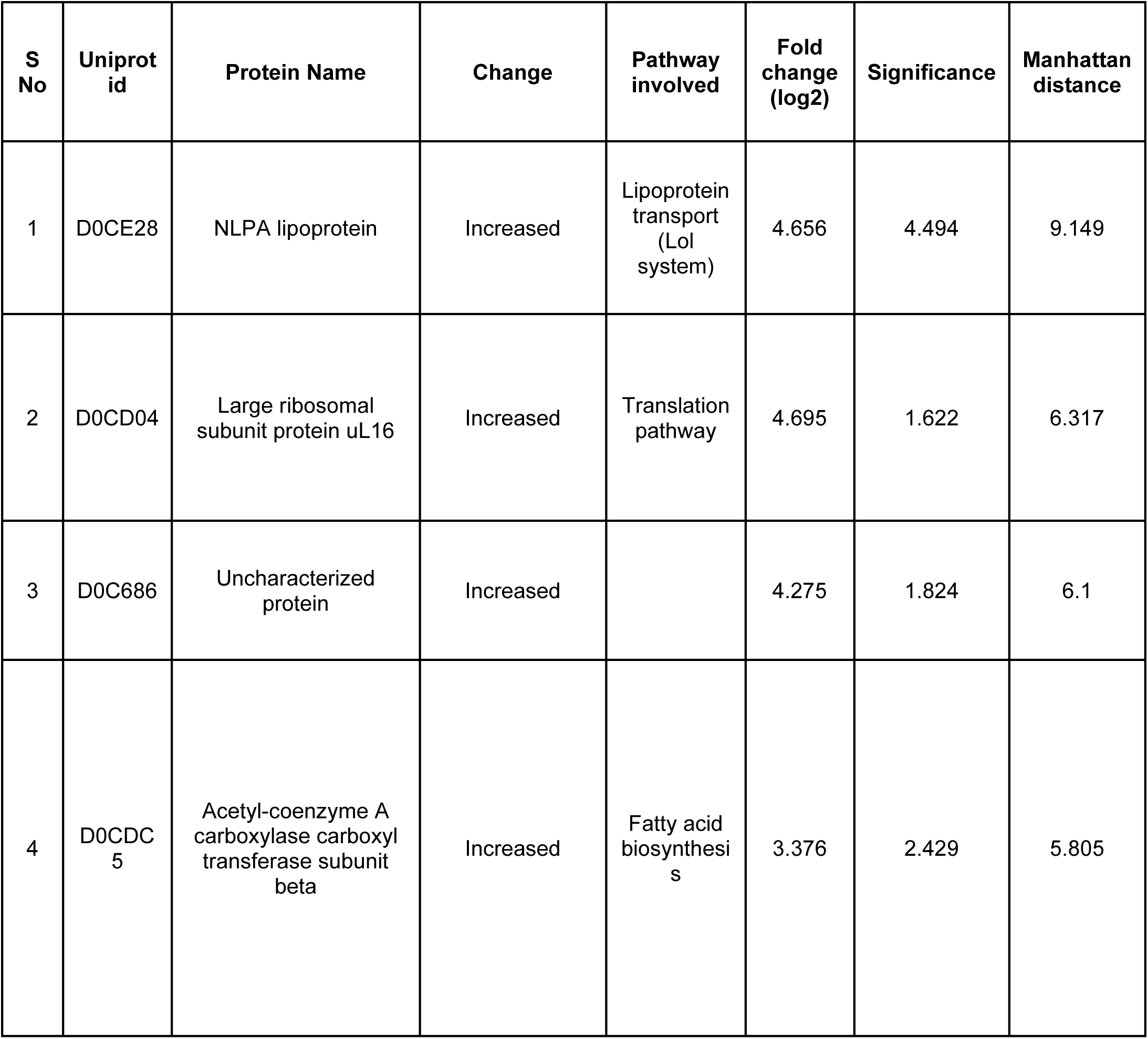
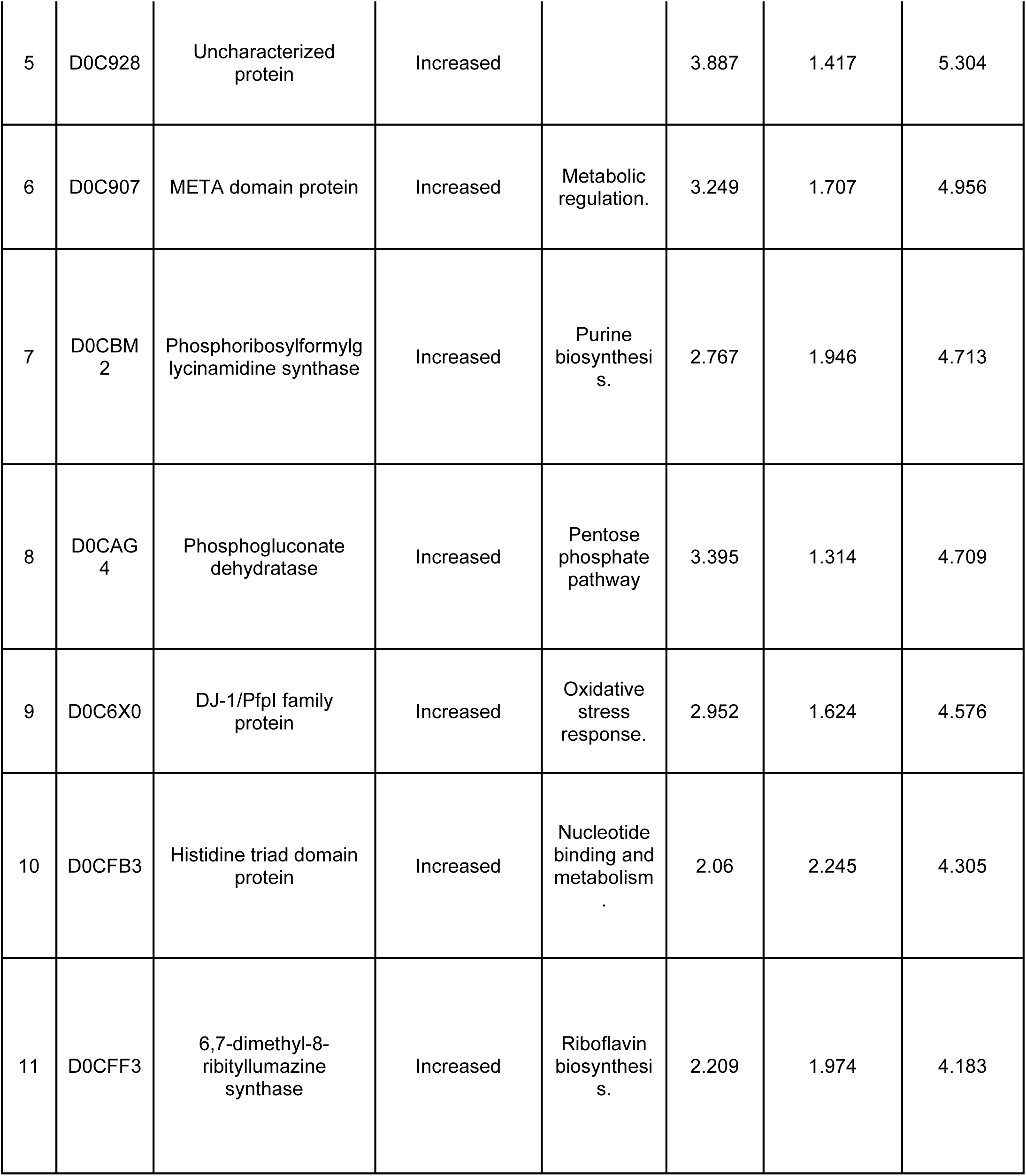
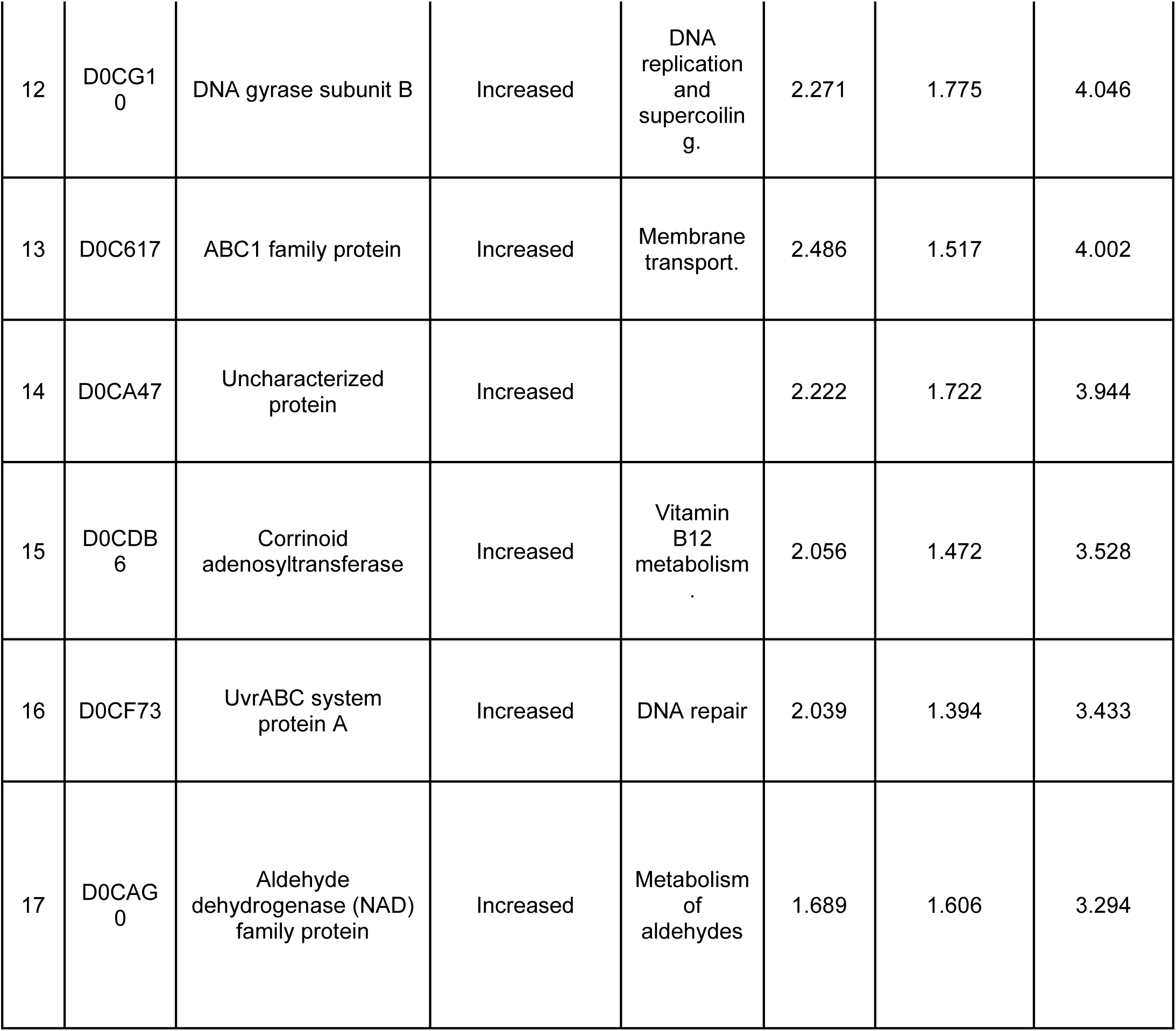
List of upregulated proteins.

##### i. Surface associated and adhesion related proteins

Of these, the NlpA lipoprotein (Uniprot ID: D0CE28) showed a 4.6-fold increase. This protein is known for facilitating the production of outer membrane vesicles (OMVs) and enhancing cellular adhesion, both of which are essential for biofilm formation, nutrient transfer, and evading the immune system. It is also involved in methionine import and may boost immune responses. Research has identified it as a potential vaccine candidate against biofilm-forming *A. baumannii* ^31^. This nonessential protein is associated with the inner membrane and was first identified in *E. coli* ^32^. In *E. coli*, NlpA is regulated by CsgD, a key transcriptional regulator involved in biofilm development, whose expression begins in the mid-logarithmic phase and persists till the stationary phase ^33^. Other upregulated surface-associated proteins include, histidine triad domain protein (UniProt ID: D0CFB3), previously reported to be associated with virulence in *Streptococcus sp*. The rapid development and advantageous selection of this protein beyond its conserved regions suggest an adaptive role in immune evasion ^34^. Similarly, the META domain protein (Uniprot ID: D0C907) identified in different bacterial species is linked to motility and virulence ^35^. Studies in *E. coli* and *Pseudomonas* revealed that adherence to softer surfaces activates essential biofilm regulators, resulting in reduced motility and increased biofilm formation ^36^. Likewise, the ABC family transporter (UniProt ID: D0C617) protein enables the translocation of multiple substrates, like metal ions and proteins, across cellular membranes by pairing this activity with ATP hydrolysis. These are involved in key processes that include multidrug resistance, nutrition absorption, adhesion, sporulation, conjugation, biofilm development, and toxin secretion. Mutagenesis studies on *E. coli, S. aureus*, and *S. pneumoniae* highlight their significance in the pathogenicity of disease ^37^.

##### ii. Stress-response and adaptive defense mechanism related

Upregulated stress-associated proteins suggest enhanced defense capabilities in strong biofilm formers. The DJ-1/PfpI family protein (Uniprot ID: D0C6X0) makes up part of the DJ-1/ThiJ/PfpI superfamily, with homologs found in several species, including humans, *Drosophila melanogaster*, *Caenorhabditis elegans*, *E. coli*, and yeast. Proteins within this family are mostly linked to activities like oxidative stress response and RNA binding. The *E. coli*, homolog referred to as Hsp31 has activities like chaperone action, acid resistance, and extensive aminopeptidase activity. While its direct participation in biofilm formation has not been shown in any bacterial species, its function in stress responses implies a possible indirect contribution to biofilm growth ^38, 39^. Additionally, the large ribosomal subunit protein uL16 (Uniprot ID: D0CD04), components of the translation machinery, was found elevated, suggesting an enhanced protein synthesis mechanism to support bacterial survival. A study reported that uL16, a crucial element of the 50S large ribosomal subunit, is essential for appropriate ribosome maturation and functionality. The lack of it may hinder the bacterium’s capacity for protein synthesis, thereby affecting biofilm stability ^40^.

##### iii. Translation-associated protein

Further, the protein A of the UvrABC system (Uniprot ID: D0CF73) is known to be involved in the activation of the adaptive defensive mechanisms of bacteria. This protein is involved in the nucleotide excision repair (NER) pathway, where UvrA and UvrB form a complex when DNA damage is observed. After UvrA has identified the DNA helix aberrations, UvrB confirms the existence of a lesion and firmly binds to the damaged site, leading to the release of UvrA ^41^. In a study on *E.coli,* uvrABC system protein A was found to be upregulated in the presence of a bioreductive agent ^42^.

##### iv. Replication and transcription machinery

The ATP-dependent enzyme DNA gyrase subunit B (Uniprot ID: D0CG10) was also found to be upregulated, which is essential for chromosomal segregation, DNA replication, and transcription. Its ubiquitous presence in all bacteria makes it a compelling and feasible target for the development of new antibacterial drugs ^43^. Prior research has emphasized DNA gyrase inhibitors from many origins as potent frameworks for drug development.

Their synthesis methodologies, structure–activity correlations, and extensive efficacy provide them exemplary prospects for the development of fresh therapies targeting resistant bacteria and biofilms ^44^. An integrated in vitro and in silico study in *Bacillus subtilis* and *Staphylococcus aureus* found marine-derived natural compounds as prospective DNA gyrase inhibitors, leading to identification of new antibacterial and anti-biofilm compounds ^45^.

##### v. Metabolic and biosynthetic enzymes

Metabolic reprogramming was evident in strong biofilm formers, with multiple biosynthetic enzymes enriched. Acetyl-CoA carboxylase beta subunit (Uniprot ID: D0CDC5) appeared prominently, indicating a rise in fatty acid biosynthesis. It is a biotin-dependent enzyme that catalyzes the carboxylation of acetyl-CoA to malonyl-CoA, a crucial step in fatty acid biosynthesis. In bacteria, it is a multi-subunit complex (accD) which forms malonyl-coA. This serves as the building block for the elongation of fatty acids and essential components of the bacterial cell membrane. Fatty acids are the precursors of LPS (Lipopolysaccharides), providing structural integrity to the biofilms ^46^. In *Mycobacterium tuberculosis,* the dense mycolate layer in the cell wall confers resistance to chemicals and desiccation while facilitating biofilm formation^47, 48^. Another biosynthetic pathway enzyme phosphoribosyl formyl glycinamidine synthase (Uniprot ID: D0CBM2) showed increased expression, indicating an upsurge in purine biosynthesis. This enzyme participates in the purine biosynthesis process. This facilitates the ATP-dependent conversion of formylglycinamide ribonucleotide (FGAR) and glutamine into formyl glycinamidine ribonucleotide (FGAM) or purL and glutamate ^49^. A study in *E. coli* identifies disrupted purL as essential for curli production and biofilm formation ^50^. Research suggests that purine metabolism affects microbe-host interactions, however, the underlying mechanisms are not clear. In bacterium *Photorhabdus temperata*, purL was identified as increased in nematode infective juveniles, and its deletion reduced both biofilm formation and bacterial persistence, underscoring its significance in symbiosis ^51^. The observed upregulation of phosphogluconate dehydratase (Uniprot ID: D0CAG4), an enzyme of the Entner-Doudoroff pathway, suggests the potential activation of non-classical sugar metabolism in gram-negative bacteria. A study indicates that this enzyme is crucial for both the use of glycerol and the enhancement of biofilm strength in its presence. This indicates that gluconeogenesis and glucose degradation influence both glycerol uses and the proliferation of glycerol-induced biofilms ^52^.

Certain enzymes have become potent antibacterial targets by interfering with important metabolic or survival processes, even if they do not directly affect the production of biofilms. This underlines their capability in formulating effective antimicrobial treatments. Like 6,7-dimethyl-8-ribityllumazine synthase (UniProt ID: D0CFF3) catalyzes the last reaction in riboflavin biosynthesis by condensing 5-amino-6-(D-ribitylamino) uracil with 3,4-dihydroxy-2-butanone 4-phosphate to produce 6,7-dimethyl-8-ribityllumazine. A study on *MtB* established RibH as a viable drug target for the development of novel antibacterial agents ^53^. Proteins belonging to the aldehyde dehydrogenase family (UniProt ID: D0CAG0) are vital enzymes that maintain bacteria’s tremendous metabolic capabilities^54^.

A restricted set of uncharacterized proteins (e.g., D0CA47, D0C686, and D0C928) exhibiting substantial fold change and importance suggests possible fresh targets that may be crucial for pathogenesis. Collectively, these findings illustrate how strong biofilm-forming *A. baumannii* orchestrates a coordinated response. The consistent enrichment of druggable enzymes such as DNA gyrase, riboflavin biosynthesis enzymes, and acetyl-CoA carboxylase further highlights potential intervention points for anti-biofilm therapeutics.

#### B. Down-regulated proteins

The proteins found to be downregulated in strong biofilm formers are listed in Table 3. The reduced levels indicate a reprioritization of bacterial processes during biofilm development, namely in translation, energy conservation, secretion systems, and metabolic pathways. (Table 2 or 3)

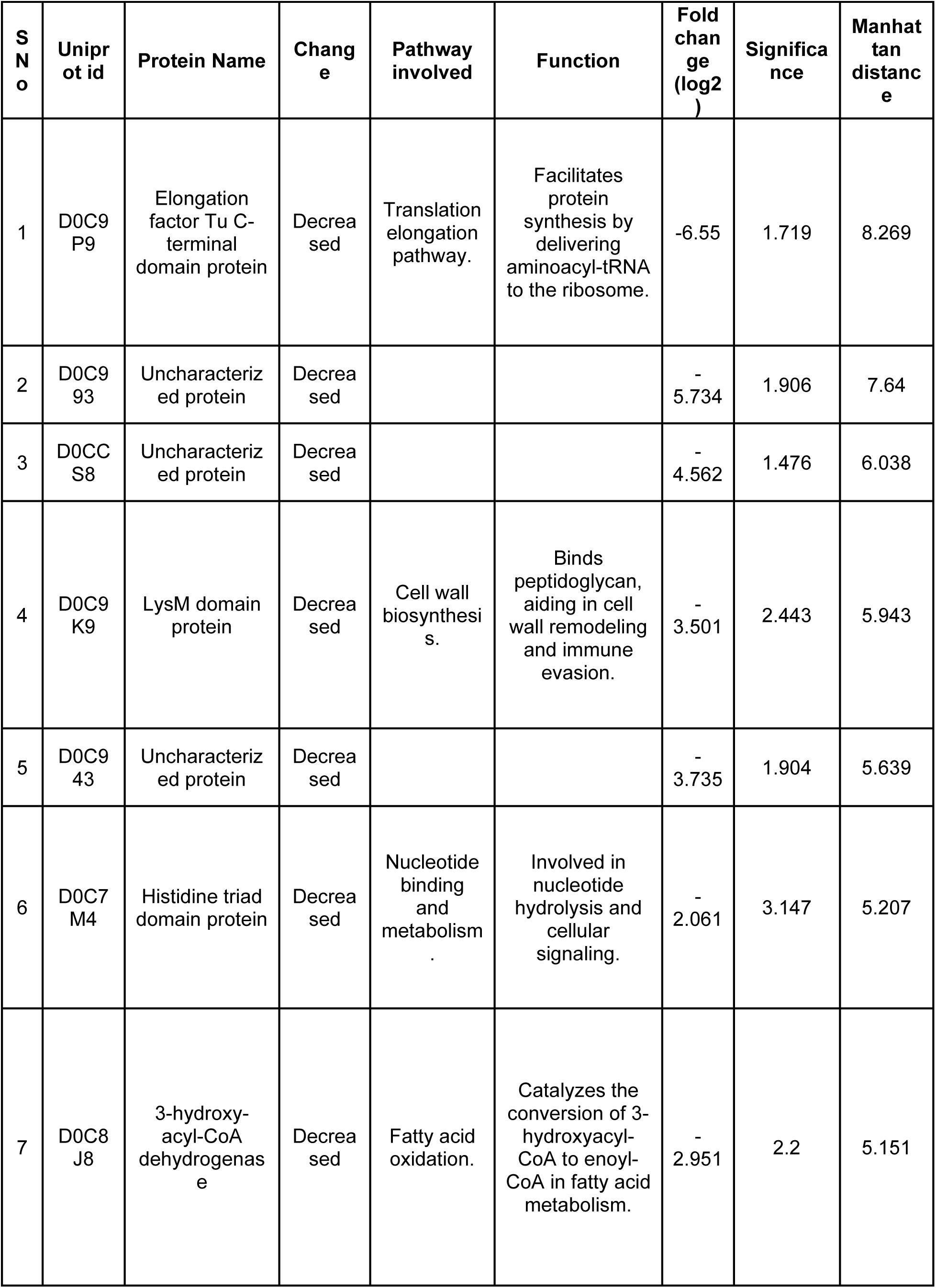
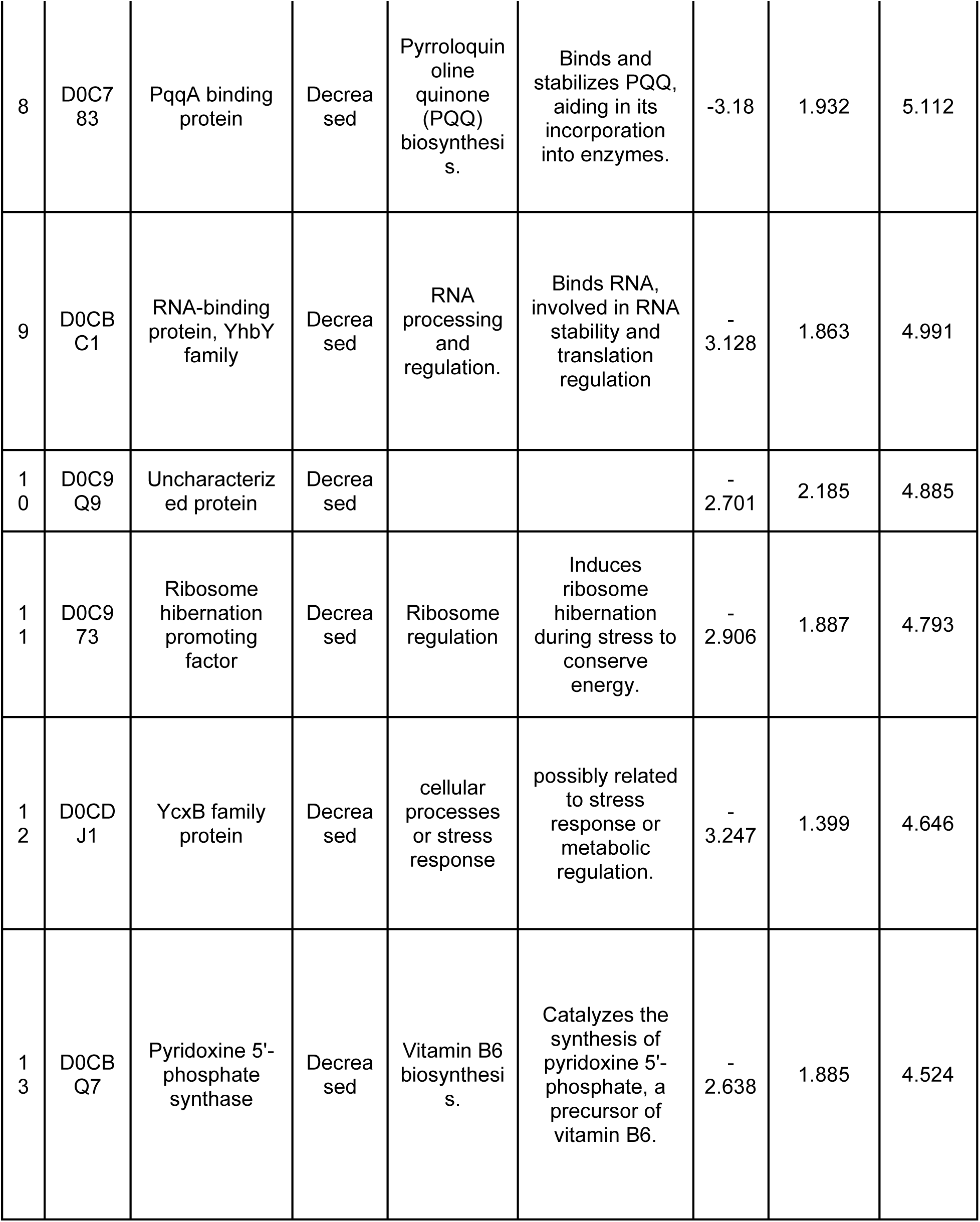
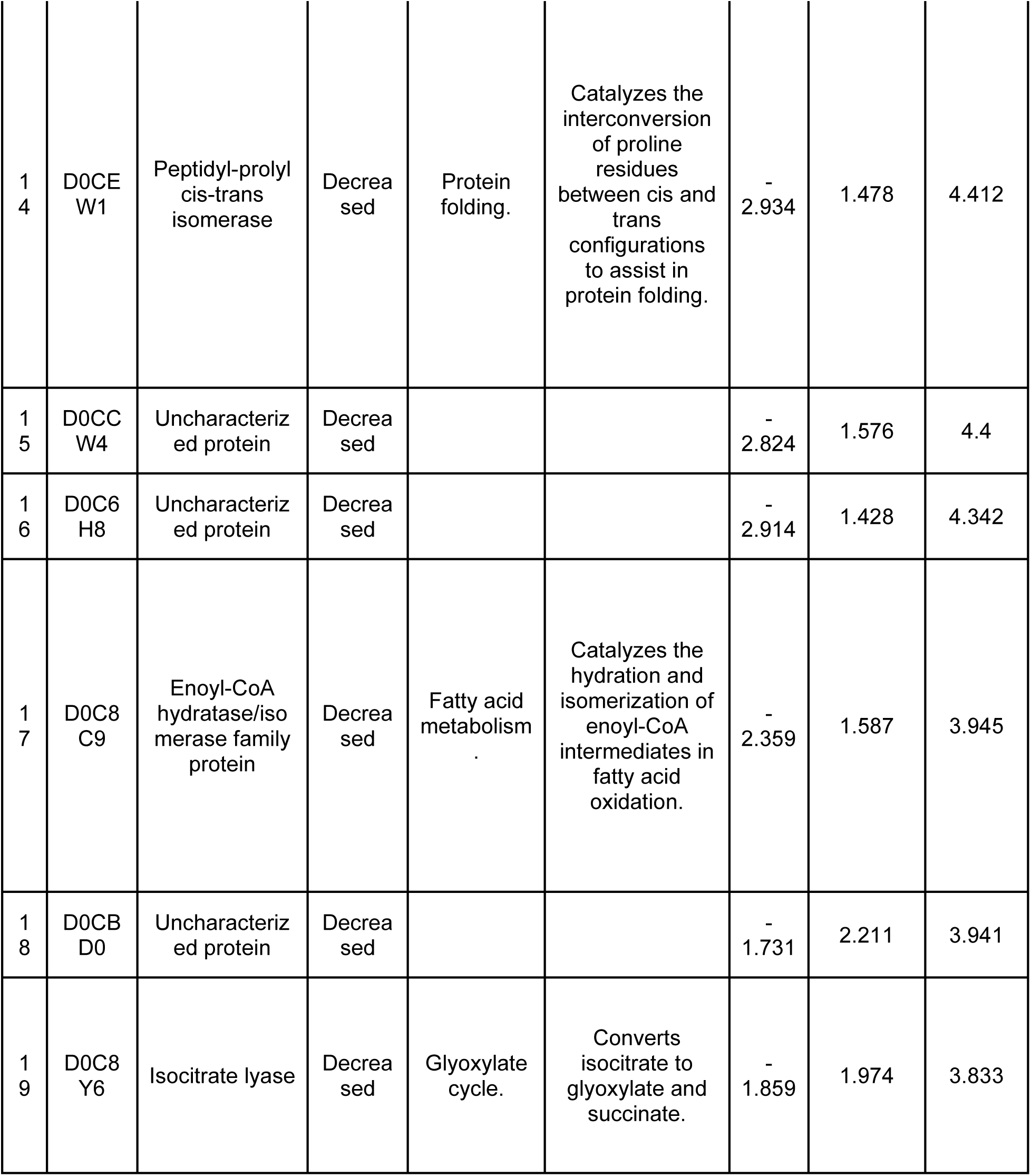
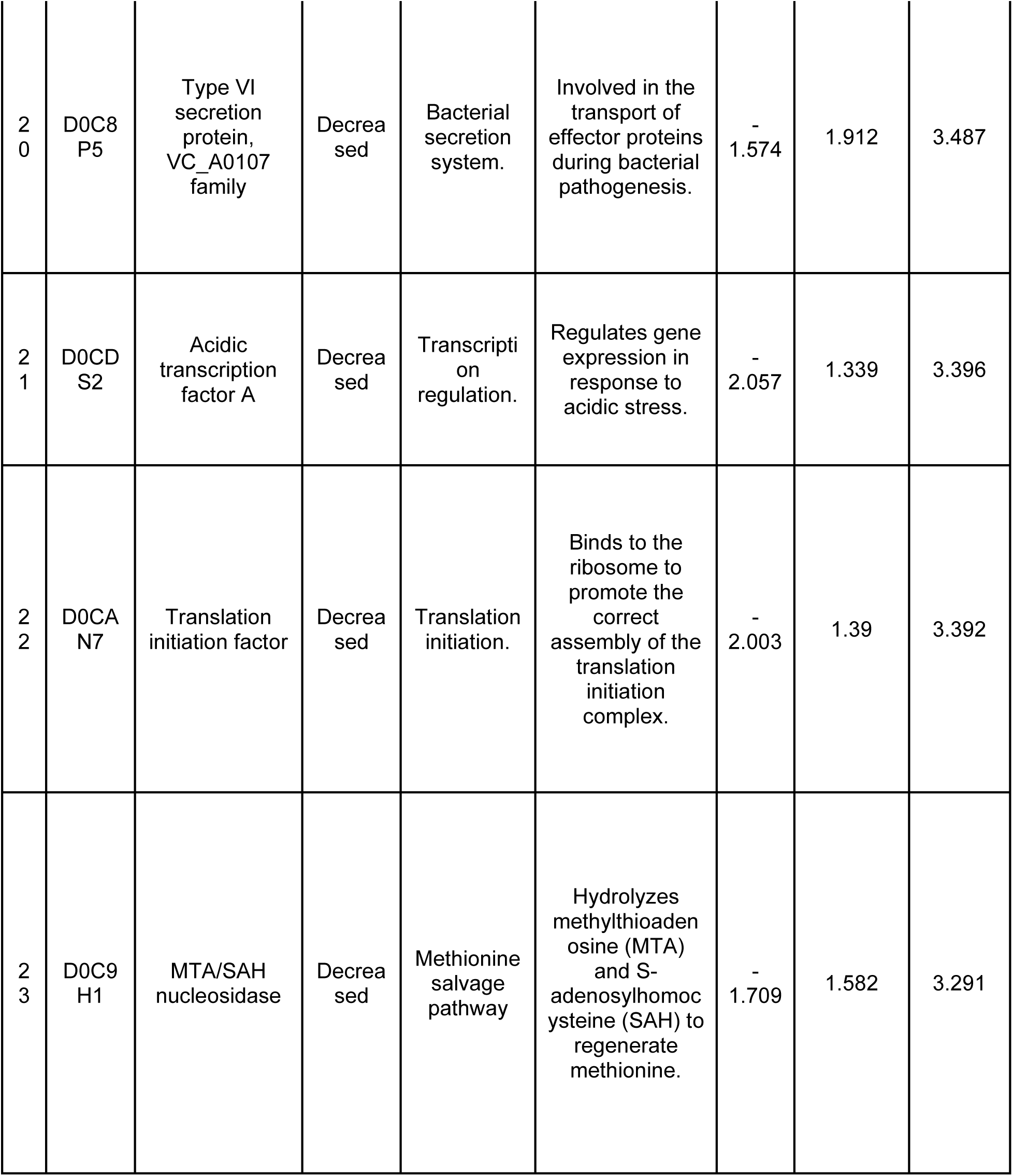
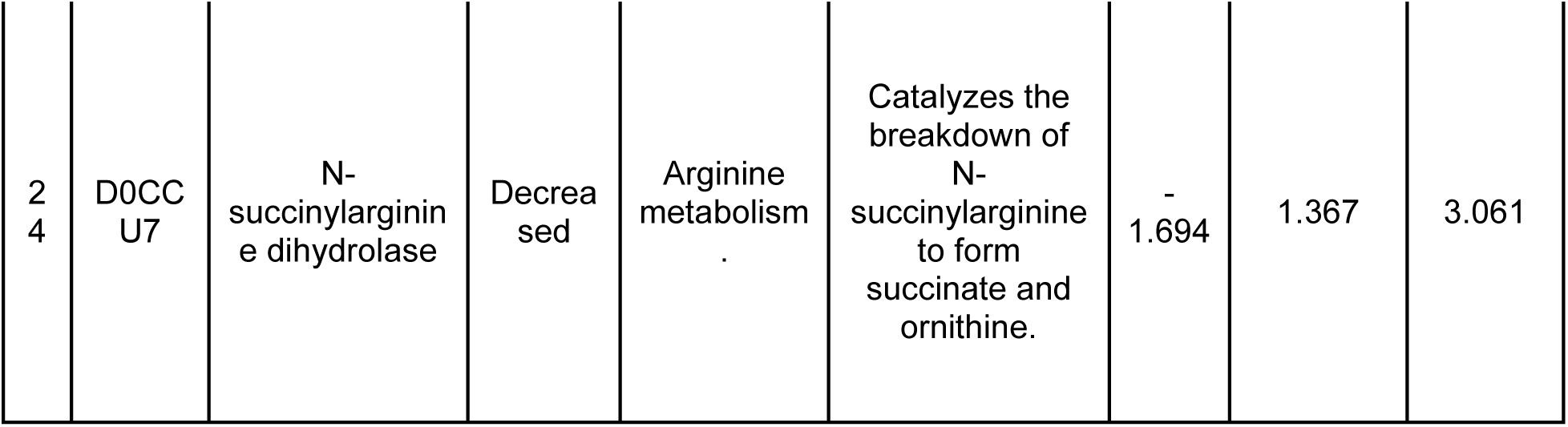

##### i. Translation factors and ribosome-associated proteins

The elongation factor thermo unstable (Ef-Tu) C-terminal domain protein (Uniprot ID: D0C9P9), central to translation elongation, showed the most significant downregulation, indicating a possible shift in protein synthesis strategy. It is one of the most abundant proteins in bacteria. It functions as an essential and universally conserved GTPase that ensures translational accuracy ^55^. Likewise, proteins linked to energy-conserving processes, like the ribosome hibernation inducing factor (Uniprot ID: D0C973), were significantly reduced, perhaps indicating a metabolic compromise under stress. *P. aeruginosa* biofilms exhibit spatial physiological heterogeneity resulting from microenvironmental adaptation. Surface-associated cells remain metabolically active, characterized by elevated ribosomal protein transcript levels, while ribosomal RNA is distributed throughout, indicating ribosome stability in dormant areas. However, deeper biofilm layers are slow-growing and are in a dormant condition expressing ribosome hibernation factors such as *rmf.* Ribosomal RNA is, however, spread throughout the biofilm, suggesting ribosome stability within the dormant regions. This stratification renders surface cells more vulnerable to antibiotics, while deeper cells exhibit increased tolerance, emphasizing spatially different roles that combined enhance biofilm resistance^56^.

##### ii. Secretion System and Virulence Factor

The type VI secretion system (T6SS) (Uniprot ID: D0C8P5), widely distributed in Gram-negative bacteria, facilitates stress responses to stimuli such as reactive oxygen species, temperature fluctuations, and pH alterations. It also plays crucial functions in bacterial growth, invasion, and virulence. The type VI secretion system (T6SS) has been discovered as a newly characterized virulence factor common in *A. baumannii*. T6SS-positive strains exhibited significantly reduced biofilm formation compared to T6SS-negative bacteria, consistent with our findings ^57^.

##### iii. Central Metabolism and related Stress-Adaptation proteins

Among metabolic enzymes, MTA/SAH nucleosidase (Uniprot ID: D0C9H1) is essential in the active methyl cycle, facilitating the recycling of adenine and methionine while aiding quorum-sensing and SAM-dependent processes. It eliminates inhibitory metabolites such as MTA and SAH. Inhibiting this enzyme hinders biofilm development and pathogenicity, making it a potential broad-spectrum antibacterial target ^58^. A study on this bacterium identified it as a homologous substrate of the type II secretion system (T2SS)^59^. Additionally, fatty acid metabolism-related enzyme 3-Hydroxyacyl-CoA dehydrogenase (Uniprot ID: D0C8J8) facilitates the third step in fatty acid β-oxidation by oxidizing the hydroxyl group of 3-hydroxyacyl-CoA to a keto group. Distinct subfamilies of this group of enzymes have been categorized according to their roles in certain metabolic processes, like fatty acid β-oxidation and amino acid degradation, as well as their substrate specificities ^60^ ^61^. Similarly, another enzyme related to vitamin biosynthesis pyridoxine 5’-phosphate synthase, (Uniprot ID: D0CBQ7), catalyzes the complex ring-closure reaction between the acyclic substrates 1-deoxy-D-xylulose-5-phosphate (DXP) and 3-amino-2-oxopropyl phosphate (1-amino-acetone-3-phosphate, AAP), yielding the formation of pyridoxine 5′-phosphate (PNP) and inorganic phosphate ^62^.

Another enzyme, Isocitrate lyase (Uniprot ID: D0C8Y6), catalyzes the cleavage of isocitrate into glyoxylate and succinate ^63^. A study revealed that these enzymes play a role in antioxidant defense, thereby contributing to antibiotic tolerance ^64^.

### 3.3 Pathway enrichment analysis of up and down-regulated proteins of biofilm phenotypes

The protein–protein interaction network demonstrated clusters of connectivity among the differentially expressed proteins. Translation-related proteins, such as *rplP* (ribosomal protein L16/uL16), *infA* (translation initiation factor IF-1), and *relA* (GTP pyrophosphokinase), constituted a densely linked hub, with chaperone *fkpA* and AGQ13237.1 (hypothetical protein). A second module was constituted by *gyrB* (DNA gyrase subunit B) and *uvrA* (UvrABC repair protein A), showing a strong association between DNA topology and repair mechanisms. Smaller clusters were identified, including *fxsA*–AGQ13440.1 and AGQ13875.1–AGQ14720.1, indicating limited functional connections (Figure 4).

**Figure 4:**
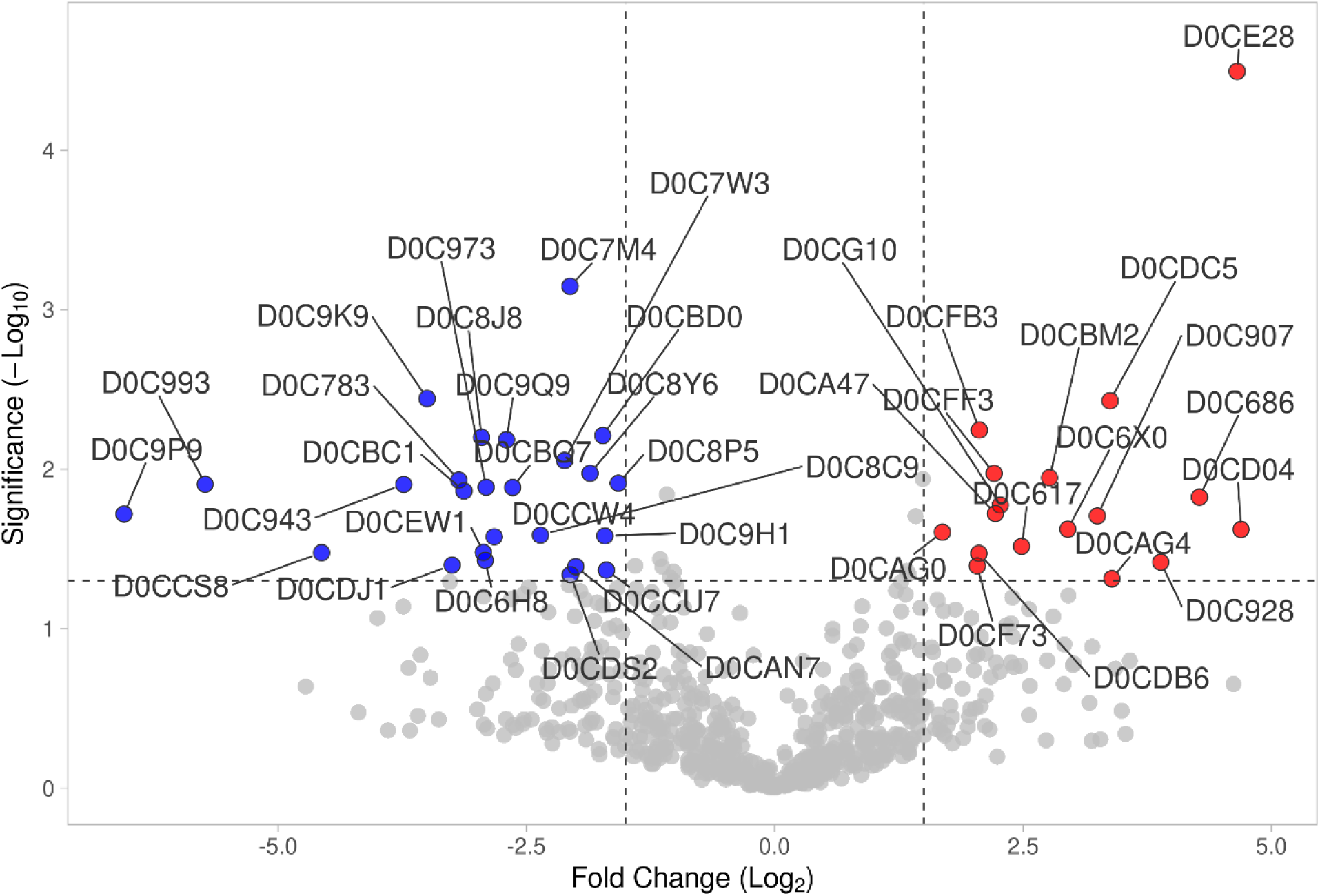
Principal Component Analysis (PCA) of *A. baumannii* clinical isolates. PCA plot showing distinct clustering of strong (red) and weak (blue) biofilm-forming isolates. Replicates from each group cluster closely together, while strong and weak biofilm formers are clearly separated along PC1 (95.9%), indicating major proteomic differences associated with biofilm phenotype.

### 3.4 Correlation of proteomics data with Quantitative Real-time PCR (qPCR)

To ensure the precision of the proteomic analysis, qPCR was used to obtain gene expression profiles of specific genes. It was performed on two additional strong biofilm-forming isolates. The genes chosen for qPCR analysis were representative of the pathways that were most significantly impacted. The majority of genes exhibiting upregulation in the proteomics dataset similarly showed substantially increased expression across all strong biofilm formers, hence confirming the reproducible nature of the results. A subset of three genes showed downregulation in these isolates, suggesting potential strain-specific variations in the regulation of particular biofilm-associated genes. Likewise, in the downregulated genes we observed substantial decreased expression, except two genes. This variability highlights the intricacy of biofilm control in *A. baumannii* and the need to examine and analyze multiple isolates to include the whole range of biofilm-related responses (Figure 5). The comparison of qPCR findings with proteomic data using a Pearson correlation matrix indicated little expression overlap between strong and weak biofilm profiles. This data substantiates the notion that proteins linked to strong biofilms operate within coordinated functional groups, while such co-regulation is absent under situations conducive to weaker biofilm formation (Figure 6). Our proteomics results point to *NlpA*, DNA gyrase B, and 6,7-dimethyl-8-ribityllumazine synthase as potential therapeutic targets.

**Figure 5:**
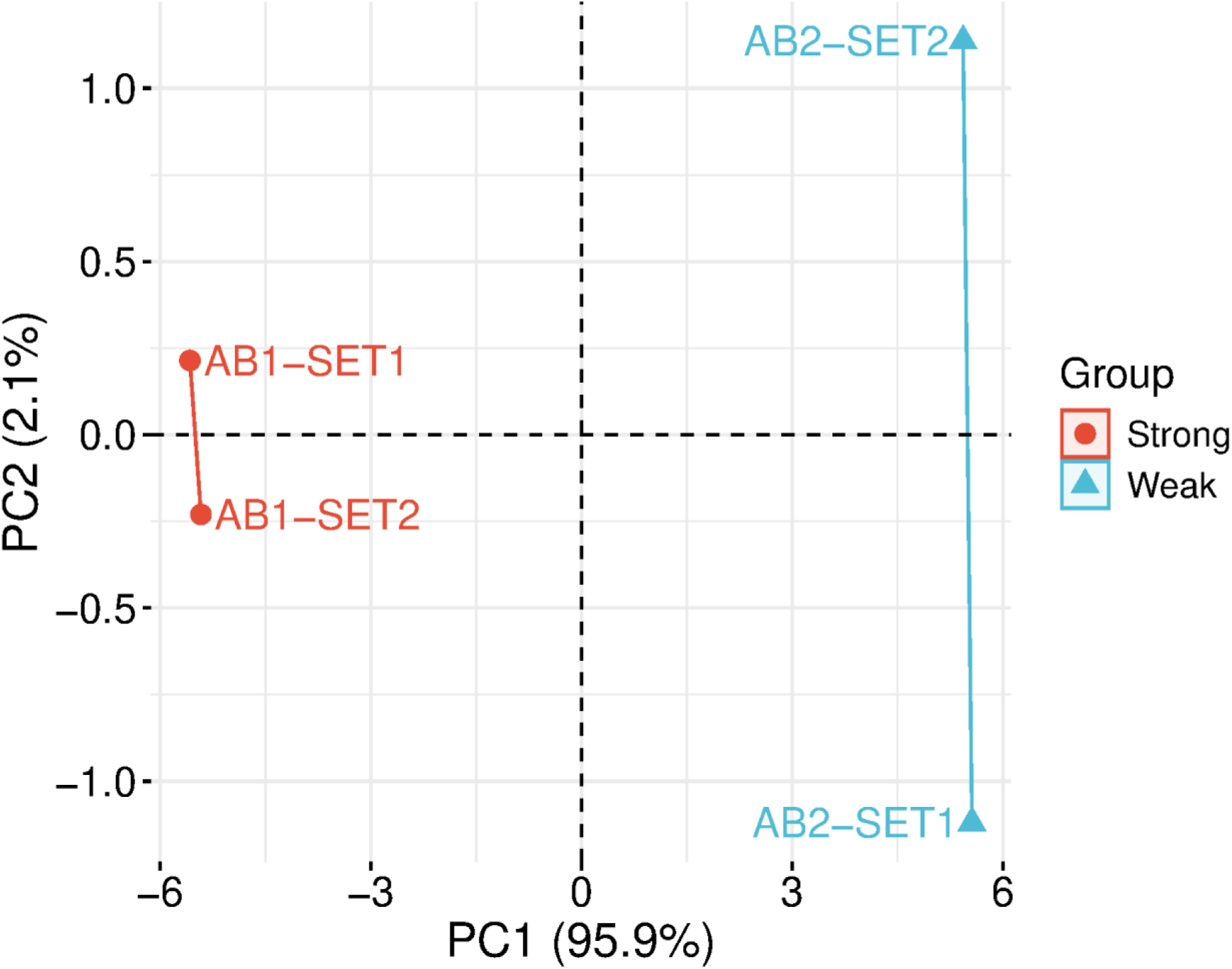
Correlation analysis of proteomic profiles from strong and weak biofilm-forming *A. baumannii* isolates. Correlation matrix showing pairwise Pearson correlation coefficients among biological replicates of strong (AB1-SET1, AB1-SET2) and weak (AB2-SET1, AB2-SET2) biofilm formers. Strong intra-group correlations (r ≈ 0.95–0.96) confirm reproducibility within each phenotype, while weak inter-group correlations (r ≈ −0.05 to −0.07) highlight clear differences between strong and weak biofilm-forming isolates.

**Figure 6A:**
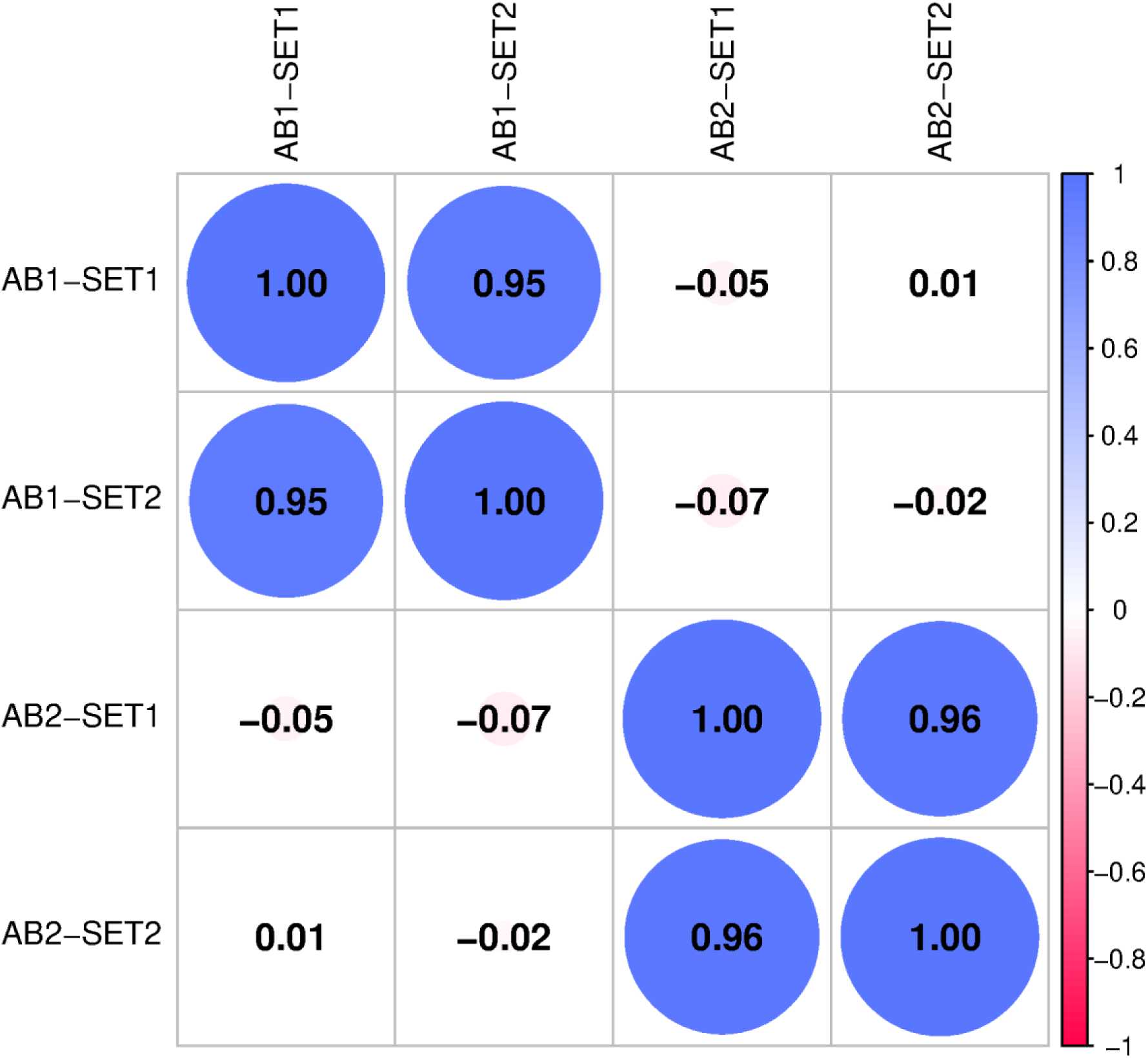
Real-time PCR validation - Upregulated proteins in strong biofilm-forming *A. baumannii* isolates compared to weak biofilm formers. Bar graph showing the log₂ fold change of significantly upregulated proteins in strong biofilm-forming isolates (AB_1, AB_2, AB_5, AB_6) relative to weak biofilm strain. Proteins involved in metabolic processes, stress response, ribosomal function, and lipoproteins were enriched, suggesting their potential roles as key biofilm determinants.

**Figure 6B:**
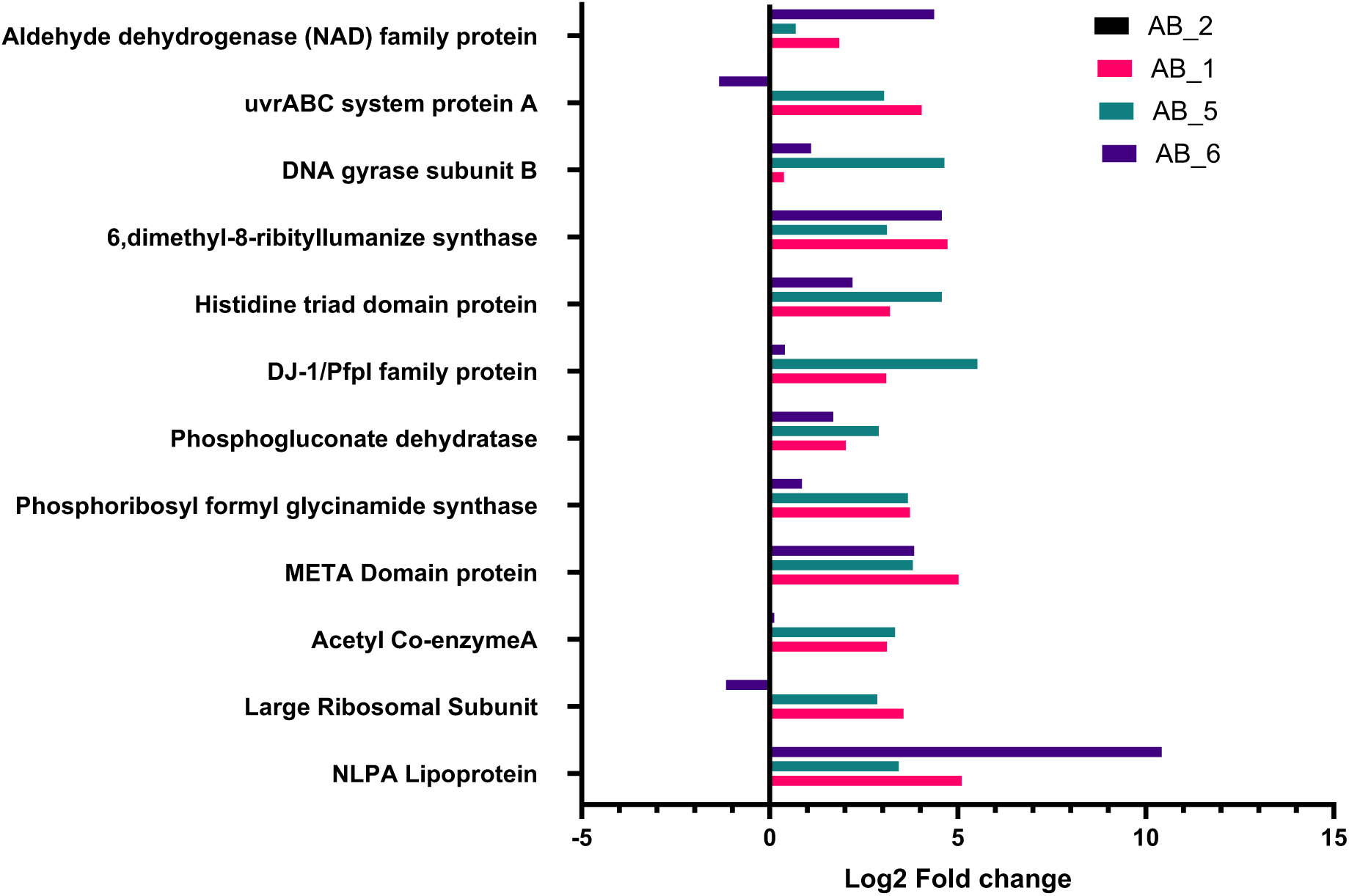
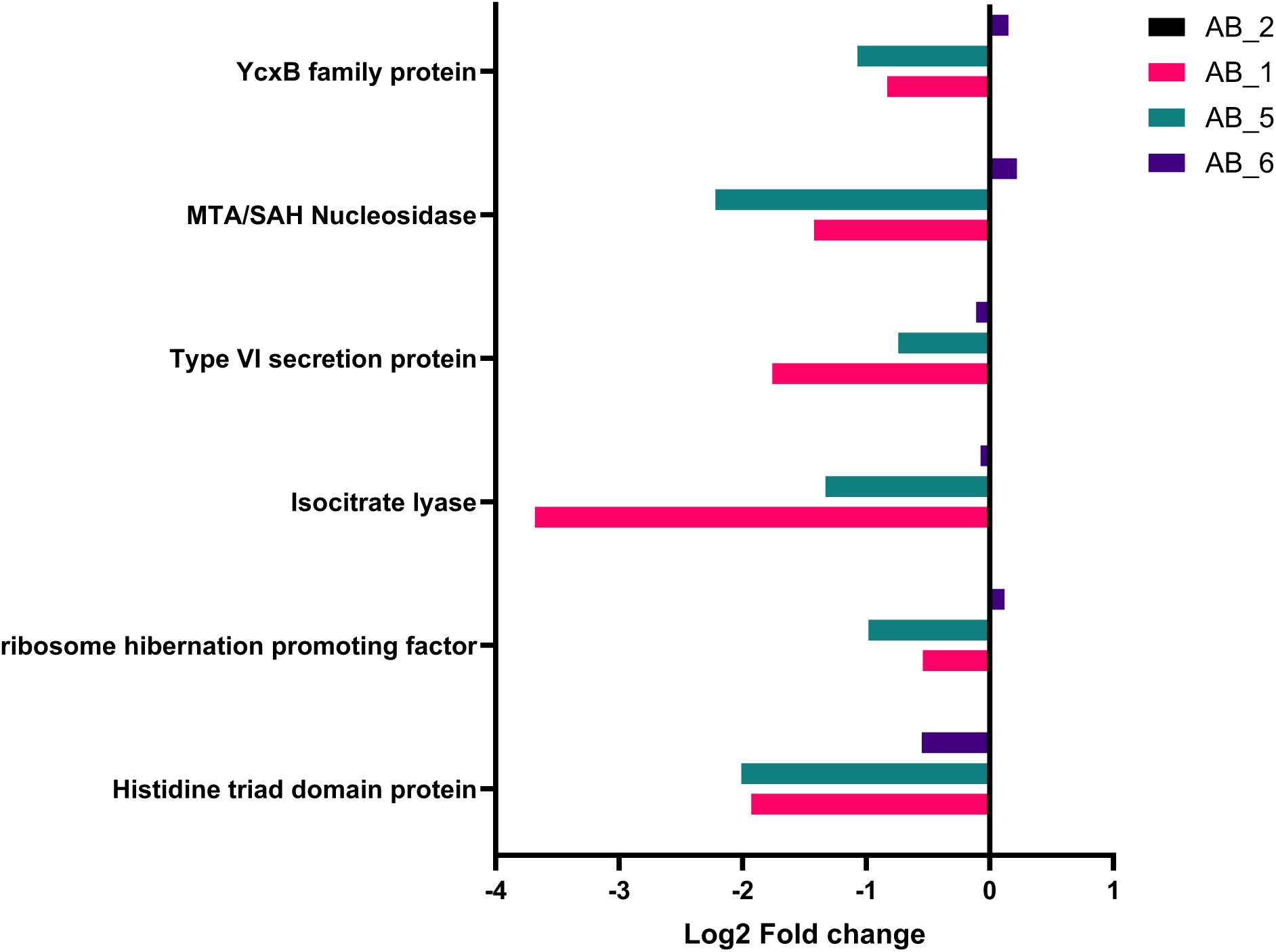
Real-time PCR validation - Downregulated proteins in strong biofilm-forming *A. baumannii* isolates compared to weak biofilm formers. Bar graph showing the log₂ fold change of significantly downregulated proteins in strong biofilm-forming isolates (AB_1, AB_2, AB_5, AB_6) relative to the weak biofilm strain. Proteins involved in translation, energy conservation, secretion systems, and metabolic pathways were enriched, suggesting their potential roles as key determinants of biofilms.

## Conclusion

By integrating quantitative protein expression profiling with statistical and functional bioinformatics analyses, this work delineates the molecular basis of biofilm robustness in *A. baumannii*. The biofilm phenotypes identify specific patterns of upregulated and downregulated proteins, highlighting the intricacy of biofilm formation and adaptive survival mechanisms. The current study presents the first comprehensive proteomic profiling among strong and weak biofilm-forming *A. baumannii* clinical isolates. Validation of the unknown regulators of biofilm formation found from this work is backed by the correlation of the acquired proteomics data with qPCR outcomes. *NlpA* lipoprotein showed promise as a target against biofilm-related infections. To confirm its role and therapeutic value, future research must include gene knockdown studies with *in-silico* inhibitor screening. These studies might lead to new approaches to combat against biofilm-forming *A. baumannii*.

## Acknowledgements

We acknowledge the support of National Institute of Pharmaceutical Education and Research, Hyderabad, and Department of Pharmaceuticals (DoP), Ministry of Chemical and Fertilizer. UB thanks ICMR-SRF grant (AMR/Fellowship/23/2022-ECD-ll) and HB thanks ICMR IIRP (2023-4907) grant for providing fellowship.

## Ethical approval

Not required

## Data availability

Not applicable

## References

1. Whiteway, C.; Breine, A.; Philippe, C.; Van der Henst, C., Acinetobacter baumannii. Trends Microbiol 2022, 30 (2), 199–200.

2. Maure, A.; Robino, E.; Van der Henst, C., The intracellular life of Acinetobacter baumannii. Trends Microbiol 2023, 31 (12), 1238–1250.

3. Miller, W. R.; Arias, C. A., ESKAPE pathogens: antimicrobial resistance, epidemiology, clinical impact and therapeutics. Nat Rev Microbiol 2024, 22 (10), 598–616.

4. Walia, K.; Madhumathi, J.; Veeraraghavan, B.; Chakrabarti, A.; Kapil, A.; Ray, P.; Singh, H.; Sistla, S.; Ohri, V. C., Establishing Antimicrobial Resistance Surveillance & Research Network in India: Journey so far. Indian J Med Res 2019, 149 (2), 164–179.

5. Gedefie, A.; Demsis, W.; Ashagrie, M.; Kassa, Y.; Tesfaye, M.; Tilahun, M.; Bisetegn, H.; Sahle, Z., Acinetobacter baumannii Biofilm Formation and Its Role in Disease Pathogenesis: A Review. Infect Drug Resist 2021, 14, 3711–3719.

6. Upmanyu, K.; Haq, Q. M. R.; Singh, R., Factors mediating Acinetobacter baumannii biofilm formation: Opportunities for developing therapeutics. Curr Res Microb Sci 2022, 3, 100131.

7. Andrea Prinzi, R. E. R., The Role of Bacterial Biofilms in Antimicrobial Resistance. American Society for Microbiology 2023.

8. Gaddy, J. A.; Tomaras, A. P.; Actis, L. A., The Acinetobacter baumannii 19606 OmpA protein plays a role in biofilm formation on abiotic surfaces and in the interaction of this pathogen with eukaryotic cells. Infect Immun 2009, 77 (8), 3150–60.

9. Zhang, P.; Hao, J.; Zhang, Y.; Su, J.; Sun, G.; Xie, J.; Hu, J.; Li, G., Understanding the clinical and molecular epidemiological characteristics of carbapenem-resistant Acinetobacter baumannii infections within intensive care units of three teaching hospitals. Ann Clin Microbiol Antimicrob 2025, 24 (1), 2.

10. Chandra, P.; V, R.; M, S.; Cs, S.; Mk, U., Multidrug-resistant Acinetobacter baumannii infections: looming threat in the Indian clinical setting. Expert Rev Anti Infect Ther 2022, 20 (5), 721–732.

11. Zhang, Y.; Cai, Y.; Zhang, B.; Zhang, Y. P. J., Spatially structured exchange of metabolites enhances bacterial survival and resilience in biofilms. Nat Commun 2024, 15 (1), 7575.

12. Rajangam, S. L.; Narasimhan, M. K., Current treatment strategies for targeting virulence factors and biofilm formation in Acinetobacter baumannii. Future Microbiol 2024, 19 (10), 941–961.

13. Hamidian, M.; Maharjan, R. P.; Farrugia, D. N.; Delgado, N. N.; Dinh, H.; Short, F. L.; Kostoulias, X.; Peleg, A. Y.; Paulsen, I. T.; Cain, A. K., Genomic and phenotypic analyses of diverse non-clinical Acinetobacter baumannii strains reveals strain-specific virulence and resistance capacity. Microb Genom 2022, 8 (2).

14. Choudhary, M.; Kaushik, S.; Kapil, A.; Shrivastava, R.; Vashistt, J., Decoding Acinetobacter baumannii biofilm dynamics and associated protein markers: proteomic and bioinformatics approach. Arch Microbiol 2022, 204 (4), 200.

15. Khemiri, A.; Jouenne, T.; Cosette, P., Proteomics dedicated to biofilmology: What have we learned from a decade of research? Med Microbiol Immunol 2016, 205 (1), 1–19.

16. Erko Stackebrandt, M. G., Nucleic Acid Techniques in Bacterial Systematics. Wiley 1991.

17. James S, L. I., CLSI Performance Standards for Antimicrobial Susceptibility Testing. CLSI Suppl. M100 2025.

18. Sarker, S. D.; Nahar, L.; Kumarasamy, Y., Microtitre plate-based antibacterial assay incorporating resazurin as an indicator of cell growth, and its application in the in vitro antibacterial screening of phytochemicals. Methods 2007, 42 (4), 321–4.

19. Magiorakos, A. P.; Srinivasan, A.; Carey, R. B.; Carmeli, Y.; Falagas, M. E.; Giske, C. G.; Harbarth, S.; Hindler, J. F.; Kahlmeter, G.; Olsson-Liljequist, B.; Paterson, D. L.; Rice, L. B.; Stelling, J.; Struelens, M. J.; Vatopoulos, A.; Weber, J. T.; Monnet, D. L., Multidrug-resistant, extensively drug-resistant and pandrug-resistant bacteria: an international expert proposal for interim standard definitions for acquired resistance. Clin Microbiol Infect 2012, 18 (3), 268–81.

20. Stepanovic, S.; Vukovic, D.; Dakic, I.; Savic, B.; Svabic-Vlahovic, M., A modified microtiter-plate test for quantification of staphylococcal biofilm formation. J Microbiol Methods 2000, 40 (2), 175–9.

21. Suriyanarayanan, T.; Qingsong, L.; Kwang, L. T.; Mun, L. Y.; Truong, T.; Seneviratne, C. J., Quantitative Proteomics of Strong and Weak Biofilm Formers of Enterococcus faecalis Reveals Novel Regulators of Biofilm Formation. Mol Cell Proteomics 2018, 17 (4), 643–654.

22. Singh, S.; Subudhi, M.; Moorthy, A. V.; Suresh, A.; Sharma, P., Ursolic acid induces apoptosis and disrupts host-parasite interactions in Theileria annulata-infected cells. Int J Parasitol Drugs Drug Resist 2025, 28, 100593.

23. Rudolph, J. D.; Cox, J., A Network Module for the Perseus Software for Computational Proteomics Facilitates Proteome Interaction Graph Analysis. J Proteome Res 2019, 18 (5), 2052–2064.

24. Khoshbakht, R.; Panahi, S.; Neshani, A.; Ghavidel, M.; Ghazvini, K., Novel approaches to overcome Colistin resistance in Acinetobacter baumannii: Exploring quorum quenching as a potential solution. Microb Pathog 2023, 182, 106264.

25. Özer B, V. C., Doğan Ö, Keske Ş, Ergönül Ö, Can F, Biofilm Formation of Acinetobacter baumannii Under in vitro and in vivo Colistin Exposure. Infect Dis Clin Microbiol 2019, 8.

26. Al-Shamiri, M. M.; Zhang, S.; Mi, P.; Liu, Y.; Xun, M.; Yang, E.; Ai, L.; Han, L.; Chen, Y., Phenotypic and genotypic characteristics of Acinetobacter baumannii enrolled in the relationship among antibiotic resistance, biofilm formation and motility. Microb Pathog 2021, 155, 104922.

27. Zeighami, H.; Valadkhani, F.; Shapouri, R.; Samadi, E.; Haghi, F., Virulence characteristics of multidrug resistant biofilm forming Acinetobacter baumannii isolated from intensive care unit patients. BMC Infect Dis 2019, 19 (1), 629.

28. Roy, S.; Chowdhury, G.; Mukhopadhyay, A. K.; Dutta, S.; Basu, S., Convergence of Biofilm Formation and Antibiotic Resistance in Acinetobacter baumannii Infection. Front Med (Lausanne) 2022, 9, 793615.

29. Farshadzadeh, Z.; Taheri, B.; Rahimi, S.; Shoja, S.; Pourhajibagher, M.; Haghighi, M. A.; Bahador, A., Growth Rate and Biofilm Formation Ability of Clinical and Laboratory-Evolved Colistin-Resistant Strains of Acinetobacter baumannii. Front Microbiol 2018, 9, 153.

30. Dafopoulou, K.; Xavier, B. B.; Hotterbeekx, A.; Janssens, L.; Lammens, C.; De, E.; Goossens, H.; Tsakris, A.; Malhotra-Kumar, S.; Pournaras, S., Colistin-Resistant Acinetobacter baumannii Clinical Strains with Deficient Biofilm Formation. Antimicrob Agents Chemother 2015, 60 (3), 1892–5.

31. Hashemzehi, R.; Doosti, A.; Kargar, M.; Jaafarinia, M., Cloning and expression of nlpA gene as DNA vaccine candidate against Acinetobacter baumannii. Mol Biol Rep 2018, 45 (4), 395–401.

32. Yamaguchi, K.; Inouye, M., Lipoprotein 28, an inner membrane protein of Escherichia coli encoded by nlpA, is not essential for growth. J Bacteriol 1988, 170 (8), 3747–9.

33. Schwechheimer, C.; Kuehn, M. J., Synthetic effect between envelope stress and lack of outer membrane vesicle production in Escherichia coli. J Bacteriol 2013, 195 (18), 4161–73.

34. Shao, Z. Q.; Zhang, Y. M.; Pan, X. Z.; Wang, B.; Chen, J. Q., Insight into the evolution of the histidine triad protein (HTP) family in Streptococcus. PLoS One 2013, 8 (3), e60116.

35. Schweppe, D. K.; Harding, C.; Chavez, J. D.; Wu, X.; Ramage, E.; Singh, P. K.; Manoil, C.; Bruce, J. E., Host-Microbe Protein Interactions during Bacterial Infection. Chem Biol 2015, 22 (11), 1521–1530.

36. Zheng, S.; Bawazir, M.; Dhall, A.; Kim, H. E.; He, L.; Heo, J.; Hwang, G., Implication of Surface Properties, Bacterial Motility, and Hydrodynamic Conditions on Bacterial Surface Sensing and Their Initial Adhesion. Front Bioeng Biotechnol 2021, 9, 643722.

37. Akhtar, A. A.; Turner, D. P., The role of bacterial ATP-binding cassette (ABC) transporters in pathogenesis and virulence: Therapeutic and vaccine potential. Microb Pathog 2022, 171, 105734.

38. Smith, N.; Wilson, M. A., Structural Biology of the DJ-1 Superfamily. Adv Exp Med Biol 2017, 1037, 5–24.

39. Lee, S. J.; Kim, S. J.; Kim, I. K.; Ko, J.; Jeong, C. S.; Kim, G. H.; Park, C.; Kang, S. O.; Suh, P. G.; Lee, H. S.; Cha, S. S., Crystal structures of human DJ-1 and Escherichia coli Hsp31, which share an evolutionarily conserved domain. J Biol Chem 2003, 278 (45), 44552–9.

40. Lauer, S. M.; Gasse, J.; Krizsan, A.; Reepmeyer, M.; Sprink, T.; Nikolay, R.; Spahn, C. M. T.; Hoffmann, R., The proline-rich antimicrobial peptide Api137 disrupts large ribosomal subunit assembly and induces misfolding. Nat Commun 2025, 16 (1), 567.

41. Malta, E.; Moolenaar, G. F.; Goosen, N., Dynamics of the UvrABC nucleotide excision repair proteins analyzed by fluorescence resonance energy transfer. Biochemistry 2007, 46 (31), 9080–8.

42. Wu, Z.; Wang, Y.; Li, L.; Zhen, S.; Du, H.; Wang, Z.; Xiao, S.; Wu, J.; Zhu, L.; Shen, J.; Wang, Z., New insights into the antimicrobial action and protective therapeutic effect of tirapazamine towards Escherichia coli-infected mice. Int J Antimicrob Agents 2023, 62 (3), 106923.

43. Jara, L. M.; Perez-Varela, M.; Corral, J.; Arch, M.; Cortes, P.; Bou, G.; Aranda, J.; Barbe, J., Novobiocin Inhibits the Antimicrobial Resistance Acquired through DNA Damage-Induced Mutagenesis in Acinetobacter baumannii. Antimicrob Agents Chemother 2016, 60 (1), 637–9.

44. Khan, T.; Sankhe, K.; Suvarna, V.; Sherje, A.; Patel, K.; Dravyakar, B., DNA gyrase inhibitors: Progress and synthesis of potent compounds as antibacterial agents. Biomed Pharmacother 2018, 103, 923–938.

45. Sayed, A. M.; Alhadrami, H. A.; El-Hawary, S. S.; Mohammed, R.; Hassan, H. M.; Rateb, M. E.; Abdelmohsen, U. R.; Bakeer, W., Discovery of Two Brominated Oxindole Alkaloids as Staphylococcal DNA Gyrase and Pyruvate Kinase Inhibitors via Inverse Virtual Screening. Microorganisms 2020, 8 (2).

46. James, E. S.; Cronan, J. E., Expression of two Escherichia coli acetyl-CoA carboxylase subunits is autoregulated. J Biol Chem 2004, 279 (4), 2520–7.

47. Pawelczyk, J.; Brzostek, A.; Kremer, L.; Dziadek, B.; Rumijowska-Galewicz, A.; Fiolka, M.; Dziadek, J., AccD6, a key carboxyltransferase essential for mycolic acid synthesis in Mycobacterium tuberculosis, is dispensable in a nonpathogenic strain. J Bacteriol 2011, 193 (24), 6960–72.

48. Reddy, M. C.; Breda, A.; Bruning, J. B.; Sherekar, M.; Valluru, S.; Thurman, C.; Ehrenfeld, H.; Sacchettini, J. C., Structure, activity, and inhibition of the Carboxyltransferase beta-subunit of acetyl coenzyme A carboxylase (AccD6) from Mycobacterium tuberculosis. Antimicrob Agents Chemother 2014, 58 (10), 6122–32.

49. Patterson, D.; Bleskan, J.; Gardiner, K.; Bowersox, J., Human phosphoribosylformylglycineamide amidotransferase (FGARAT): regional mapping, complete coding sequence, isolation of a functional genomic clone, and DNA sequence analysis. Gene 1999, 239 (2), 381–91.

50. Cepas, V.; Ballen, V.; Gabasa, Y.; Ramirez, M.; Lopez, Y.; Soto, S. M., Transposon Insertion in the purL Gene Induces Biofilm Depletion in Escherichia coli ATCC 25922. Pathogens 2020, 9 (9).

51. An, R.; Grewal, P. S., purL gene expression affects biofilm formation and symbiotic persistence of Photorhabdus temperata in the nematode Heterorhabditis bacteriophora. Microbiology (Reading) 2011, 157 (Pt 9), 2595–2603.

52. Pan, S.; Underhill, S. A. M.; Hamm, C. W.; Stover, M. A.; Butler, D. R.; Shults, C. A.; Manjarrez, J. R.; Cabeen, M. T., Glycerol metabolism impacts biofilm phenotypes and virulence in Pseudomonas aeruginosa via the Entner-Doudoroff pathway. mSphere 2024, 9 (4), e0078623.

53. Singh, M.; Dhanwal, A.; Verma, A.; Augustin, L.; Kumari, N.; Chakraborti, S.; Agarwal, N.; Sriram, D.; Dey, R. J., Discovery of potent antimycobacterial agents targeting lumazine synthase (RibH) of Mycobacterium tuberculosis. Sci Rep 2024, 14 (1), 12170.

54. Riveros-Rosas, H.; Julian-Sanchez, A.; Moreno-Hagelsieb, G.; Munoz-Clares, R. A., Aldehyde dehydrogenase diversity in bacteria of the Pseudomonas genus. Chem Biol Interact 2019, 304, 83–87.

55. Widjaja, M.; Harvey, K. L.; Hagemann, L.; Berry, I. J.; Jarocki, V. M.; Raymond, B. B. A.; Tacchi, J. L.; Grundel, A.; Steele, J. R.; Padula, M. P.; Charles, I. G.; Dumke, R.; Djordjevic, S. P., Elongation factor Tu is a multifunctional and processed moonlighting protein. Sci Rep 2017, 7 (1), 11227.

56. Williamson, K. S.; Richards, L. A.; Perez-Osorio, A. C.; Pitts, B.; McInnerney, K.; Stewart, P. S.; Franklin, M. J., Heterogeneity in Pseudomonas aeruginosa biofilms includes expression of ribosome hibernation factors in the antibiotic-tolerant subpopulation and hypoxia-induced stress response in the metabolically active population. J Bacteriol 2012, 194 (8), 2062–73.

57. Lin, Y.; Zhao, D.; Huang, N.; Liu, S.; Zheng, J.; Cao, J.; Zeng, W.; Zheng, X.; Wang, L.; Zhou, T.; Sun, Y., Clinical impact of the type VI secretion system on clinical characteristics, virulence and prognosis of Acinetobacter baumannii during bloodstream infection. Microb Pathog 2023, 182, 106252.

58. Parveen, N.; Cornell, K. A., Methylthioadenosine/S-adenosylhomocysteine nucleosidase, a critical enzyme for bacterial metabolism. Mol Microbiol 2011, 79 (1), 7–20.

59. Elhosseiny, N. M.; Elhezawy, N. B.; Attia, A. S., Comparative proteomics analyses of Acinetobacter baumannii strains ATCC 17978 and AB5075 reveal the differential role of type II secretion system secretomes in lung colonization and ciprofloxacin resistance. Microb Pathog 2019, 128, 20–27.

60. Xu, Y.; Sun, F., Purification, crystallization and preliminary crystallographic analysis of 3-hydroxyacyl-CoA dehydrogenase from Caenorhabditis elegans. Acta Crystallogr Sect F Struct Biol Cryst Commun 2013, 69 (Pt 5), 515–9.

61. Jimenez-Diaz, L., Caballero, A., Segura, A., Pathways for the Degradation of Fatty Acids in Bacteria. Rojo, F. (eds) Aerobic Utilization of Hydrocarbons, Oils and Lipids. Handbook of Hydrocarbon and Lipid Microbiology 2017, 23.

62. Denise, R.; Babor, J.; Gerlt, J. A.; de Crecy-Lagard, V., Pyridoxal 5’-phosphate synthesis and salvage in Bacteria and Archaea: predicting pathway variant distributions and holes. Microb Genom 2023, 9 (2).

63. Dunn, M. F.; Ramirez-Trujillo, J. A.; Hernandez-Lucas, I., Major roles of isocitrate lyase and malate synthase in bacterial and fungal pathogenesis. Microbiology (Reading) 2009, 155 (Pt 10), 3166–3175.

64. Nandakumar, M.; Nathan, C.; Rhee, K. Y., Isocitrate lyase mediates broad antibiotic tolerance in Mycobacterium tuberculosis. Nat Commun 2014, 5, 4306.

