## Supplementary Tables for "Proteomic Insights into Strong and Weak Biofilm Formation in *Acinetobacter baumannii* for Potential Therapeutic Targets"

Supplementary Table 1

| **S No** | **Uniprot id** | **Protein Name** | **Change** | **Pathway involved** | **Fold change (log2)** | **Significance** | **Manhattan distance** |
| --- | --- | --- | --- | --- | --- | --- | --- |
| 1 | D0CE28 | NLPA lipoprotein | Increased | Lipoprotein transport (Lol system) | 4.656 | 4.494 | 9.149 |
| 2 | D0CD04 | Large ribosomal subunit protein uL16 | Increased | Translation pathway | 4.695 | 1.622 | 6.317 |
| 3 | D0C686 | Uncharacterized protein | Increased |  | 4.275 | 1.824 | 6.1 |
| 4 | D0CDC5 | Acetyl-coenzyme A carboxylase carboxyl transferase subunit beta | Increased | Fatty acid biosynthesis | 3.376 | 2.429 | 5.805 |
| 5 | D0C928 | Uncharacterized protein | Increased |  | 3.887 | 1.417 | 5.304 |
| 6 | D0C907 | META domain protein | Increased | Metabolic regulation. | 3.249 | 1.707 | 4.956 |
| 7 | D0CBM2 | Phosphoribosylformylglycinamidine synthase | Increased | Purine biosynthesis. | 2.767 | 1.946 | 4.713 |
| 8 | D0CAG4 | Phosphogluconate dehydratase | Increased | Pentose phosphate pathway | 3.395 | 1.314 | 4.709 |
| 9 | D0C6X0 | DJ-1/PfpI family protein | Increased | Oxidative stress response. | 2.952 | 1.624 | 4.576 |
| 10 | D0CFB3 | Histidine triad domain protein | Increased | Nucleotide binding and metabolism. | 2.06 | 2.245 | 4.305 |
| 11 | D0CFF3 | 6,7-dimethyl-8-ribityllumazine synthase | Increased | Riboflavin biosynthesis. | 2.209 | 1.974 | 4.183 |
| 12 | D0CG10 | DNA gyrase subunit B | Increased | DNA replication and supercoiling. | 2.271 | 1.775 | 4.046 |
| 13 | D0C617 | ABC1 family protein | Increased | Membrane transport. | 2.486 | 1.517 | 4.002 |
| 14 | D0CA47 | Uncharacterized protein | Increased |  | 2.222 | 1.722 | 3.944 |
| 15 | D0CDB6 | Corrinoid adenosyltransferase | Increased | Vitamin B12 metabolism. | 2.056 | 1.472 | 3.528 |
| 16 | D0CF73 | UvrABC system protein A | Increased | DNA repair | 2.039 | 1.394 | 3.433 |
| 17 | D0CAG0 | Aldehyde dehydrogenase (NAD) family protein | Increased | Metabolism of aldehydes | 1.689 | 1.606 | 3.294 |

Supplementary Table 2

| **S No** | **Uniprot id** | **Protein Name** | **Change** | **Pathway involved** | **Function** | **Fold change (log2)** | **Significance** | **Manhattan distance** |
| --- | --- | --- | --- | --- | --- | --- | --- | --- |
| 1 | D0C9P9 | Elongation factor Tu C-terminal domain protein | Decreased | Translation elongation pathway. | Facilitates protein synthesis by delivering aminoacyl-tRNA to the ribosome. | -6.55 | 1.719 | 8.269 |
| 2 | D0C993 | Uncharacterized protein | Decreased |  |  | -5.734 | 1.906 | 7.64 |
| 3 | D0CCS8 | Uncharacterized protein | Decreased |  |  | -4.562 | 1.476 | 6.038 |
| 4 | D0C9K9 | LysM domain protein | Decreased | Cell wall biosynthesis. | Binds peptidoglycan, aiding in cell wall remodeling and immune evasion. | -3.501 | 2.443 | 5.943 |
| 5 | D0C943 | Uncharacterized protein | Decreased |  |  | -3.735 | 1.904 | 5.639 |
| 6 | D0C7M4 | Histidine triad domain protein | Decreased | Nucleotide binding and metabolism. | Involved in nucleotide hydrolysis and cellular signaling. | -2.061 | 3.147 | 5.207 |
| 7 | D0C8J8 | 3-hydroxy-acyl-CoA dehydrogenase | Decreased | Fatty acid oxidation. | Catalyzes the conversion of 3-hydroxyacyl-CoA to enoyl-CoA in fatty acid metabolism. | -2.951 | 2.2 | 5.151 |
| 8 | D0C783 | PqqA binding protein | Decreased | Pyrroloquinoline quinone (PQQ) biosynthesis. | Binds and stabilizes PQQ, aiding in its incorporation into enzymes. | -3.18 | 1.932 | 5.112 |
| 9 | D0CBC1 | RNA-binding protein, YhbY family | Decreased | RNA processing and regulation. | Binds RNA, involved in RNA stability and translation regulation | -3.128 | 1.863 | 4.991 |
| 10 | D0C9Q9 | Uncharacterized protein | Decreased |  |  | -2.701 | 2.185 | 4.885 |
| 11 | D0C973 | Ribosome hibernation promoting factor | Decreased | Ribosome regulation | Induces ribosome hibernation during stress to conserve energy. | -2.906 | 1.887 | 4.793 |
| 12 | D0CDJ1 | YcxB family protein | Decreased | cellular processes or stress response | possibly related to stress response or metabolic regulation. | -3.247 | 1.399 | 4.646 |
| 13 | D0CBQ7 | Pyridoxine 5'-phosphate synthase | Decreased | Vitamin B6 biosynthesis. | Catalyzes the synthesis of pyridoxine 5'-phosphate, a precursor of vitamin B6. | -2.638 | 1.885 | 4.524 |
| 14 | D0CEW1 | Peptidyl-prolyl cis-trans isomerase | Decreased | Protein folding. | Catalyzes the interconversion of proline residues between cis and trans configurations to assist in protein folding. | -2.934 | 1.478 | 4.412 |
| 15 | D0CCW4 | Uncharacterized protein | Decreased |  |  | -2.824 | 1.576 | 4.4 |
| 16 | D0C6H8 | Uncharacterized protein | Decreased |  |  | -2.914 | 1.428 | 4.342 |
| 17 | D0C8C9 | Enoyl-CoA hydratase/isomerase family protein | Decreased | Fatty acid metabolism. | Catalyzes the hydration and isomerization of enoyl-CoA intermediates in fatty acid oxidation. | -2.359 | 1.587 | 3.945 |
| 18 | D0CBD0 | Uncharacterized protein | Decreased |  |  | -1.731 | 2.211 | 3.941 |
| 19 | D0C8Y6 | Isocitrate lyase | Decreased | Glyoxylate cycle. | Converts isocitrate to glyoxylate and succinate. | -1.859 | 1.974 | 3.833 |
| 20 | D0C8P5 | Type VI secretion protein, VC_A0107 family | Decreased | Bacterial secretion system. | Involved in the transport of effector proteins during bacterial pathogenesis. | -1.574 | 1.912 | 3.487 |
| 21 | D0CDS2 | Acidic transcription factor A | Decreased | Transcription regulation. | Regulates gene expression in response to acidic stress. | -2.057 | 1.339 | 3.396 |
| 22 | D0CAN7 | Translation initiation factor | Decreased | Translation initiation. | Binds to the ribosome to promote the correct assembly of the translation initiation complex. | -2.003 | 1.39 | 3.392 |
| 23 | D0C9H1 | MTA/SAH nucleosidase | Decreased | Methionine salvage pathway | Hydrolyzes methylthioadenosine (MTA) and S-adenosylhomocysteine (SAH) to regenerate methionine. | -1.709 | 1.582 | 3.291 |
| 24 | D0CCU7 | N-succinylarginine dihydrolase | Decreased | Arginine metabolism. | Catalyzes the breakdown of N-succinylarginine to form succinate and ornithine. | -1.694 | 1.367 | 3.061 |
